# The Late Positive Component in Recognition Memory is Linked to Mnemonic Evidence Accumulation

**DOI:** 10.1101/2025.07.20.665783

**Authors:** Jie Sun, Daniel Feuerriegel, Mia Nightingale, Adam F. Osth

## Abstract

Recognising objects from memory requires an integration of sensory and mnemonic information. This process has been theorised to occur via a stochastic evidence accumulation process implemented within the parietal cortex. Recent electroencephalographic (EEG) evidence indicates that the widely studied parietal Late Positive Component (LPC) shows characteristics of such a mnemonic accumulation signal. Here, we formally investigated this hypothesis using generative computational modelling that links a trial-level memory strength variable in the Diffusion Decision Model (DDM) with LPC amplitudes prior to the time of the decision. We recorded EEG from 24 participants making recognition judgements based on either studied or novel words. Each participant completed up to three testing sessions. We replicated recent findings that the LPC ramps up and peaks around the time of the decision and corresponding motor response. LPC amplitudes also covaried with accuracy and response time, as expected of a neural correlate of memory strength. By fitting DDMs to LPC amplitudes and behavioural data using specialised neural network tools, we demonstrate that LPC amplitudes are selectively associated with the rate of evidence accumulation, signifying memory strength. This association was stronger for previously studied words compared to novel words, and strongest at the time window immediately prior to the recognition decision. Our findings therefore recast the LPC as a neural signature of mnemonic strength linked to the rate of evidence accumulation during recognition memory decisions.

## 1. Introduction

Decades of research using electroencephalography (EEG) recorded during recognition memory tasks have investigated the EEG correlates of memory retrieval by comparing event-related potentials (ERPs) following recognized items (targets) and novel items (lures). One extensively studied correlate, the Late Positive Component (LPC), has been consistently observed at parietal electrodes (Friedman & Johnson, 2000; Johnson et al., 1985; Kwon et al., 2023). The LPC has been widely assumed to reflect a recollection process supporting the retrieval of detailed episodic information (Rugg & Curran, 2007; Rugg & Yonelinas, 2003; Woodruff et al., 2006). LPC amplitudes are more positive-going for previously-seen compared to novel items (termed the old/new effect; Rugg & Curran, 2007), decisions made with high confidence (Addante et al., 2012; Sarah & Rugg, 2010), self-reported recollection (Nie et al., 2021; Voss & Paller, 2009), and accurate source memory judgements (Leynes & Mok, 2017; Woroch & Gonsalves, 2010). However, despite many endorsements of the LPC as a correlate of recollection, it has not been formally tested within a mechanistic framework or linked to latent variables in computational models of recognition memory.

An alternative, plausible view of the LPC is derived from the mnemonic accumulator hypothesis relating to the parietal lobe, a region of brain that was shown to be critical to episodic memory and retrieval of event details (Dobbins et al., 2003;Vilberg & Rugg, 2008; Wagner et al., 2005; Wheeler & Buckner, 2004). According to this hypothesis, parietal lobe neural activity during memory retrieval reflects the integration of mnemonic information from distributed brain regions, with the mnemonic evidence accumulating continuously over time (Donaldson et al., 2010; Sestieri et al., 2017; Wagner et al., 2005). Consistent with this view, blood oxygenation level dependent (BOLD) signals in parietal regions were found to correlate with the likelihood of items being recognised as ‘old’ (Henson et al., 1999; Hutchinson et al., 2015). Notably, the LPC has also been linked to BOLD signals and lesions in the lateral parietal cortex (Ally et al., 2008; Vilberg & Rugg, 2009). Similar to LPC amplitudes, BOLD signals in left parietal regions are greater for ‘old’ than ‘new’ judgments and for retrieval of memory details (Kahn et al., 2004; Sestieri et al., 2014; Wheeler & Buckner, 2003; Yu et al., 2012).

This mnemonic accumulation framework is often analogously compared to the classic Diffusion Decision Model (DDM; Ratcliff, 1978). This model describes mnemonic evidence accumulating via a stochastic diffusion process toward ‘old’ or ‘new’ decision boundaries. The evidence accumulates at an average speed represented by a drift rate parameter (Fig. 1a), and the model simultaneously accounts for choices and response times (RTs). Here, drift rate represents the quality or strength of the mnemonic evidence. The DDM has yielded novel insights into performance during recognition memory tasks (see Osth et al. 2025, for a review). For instance, estimates of across-trial drift rate variability have been observed to be greater for targets than lures (Starns & Ratcliff, 2014), consistent with findings relating to underlying memory strength distributions (Chen et al., 2024; Osth et al., 2017; Ratcliff et al., 1994; Wixted, 2007).

**Figure 1.**
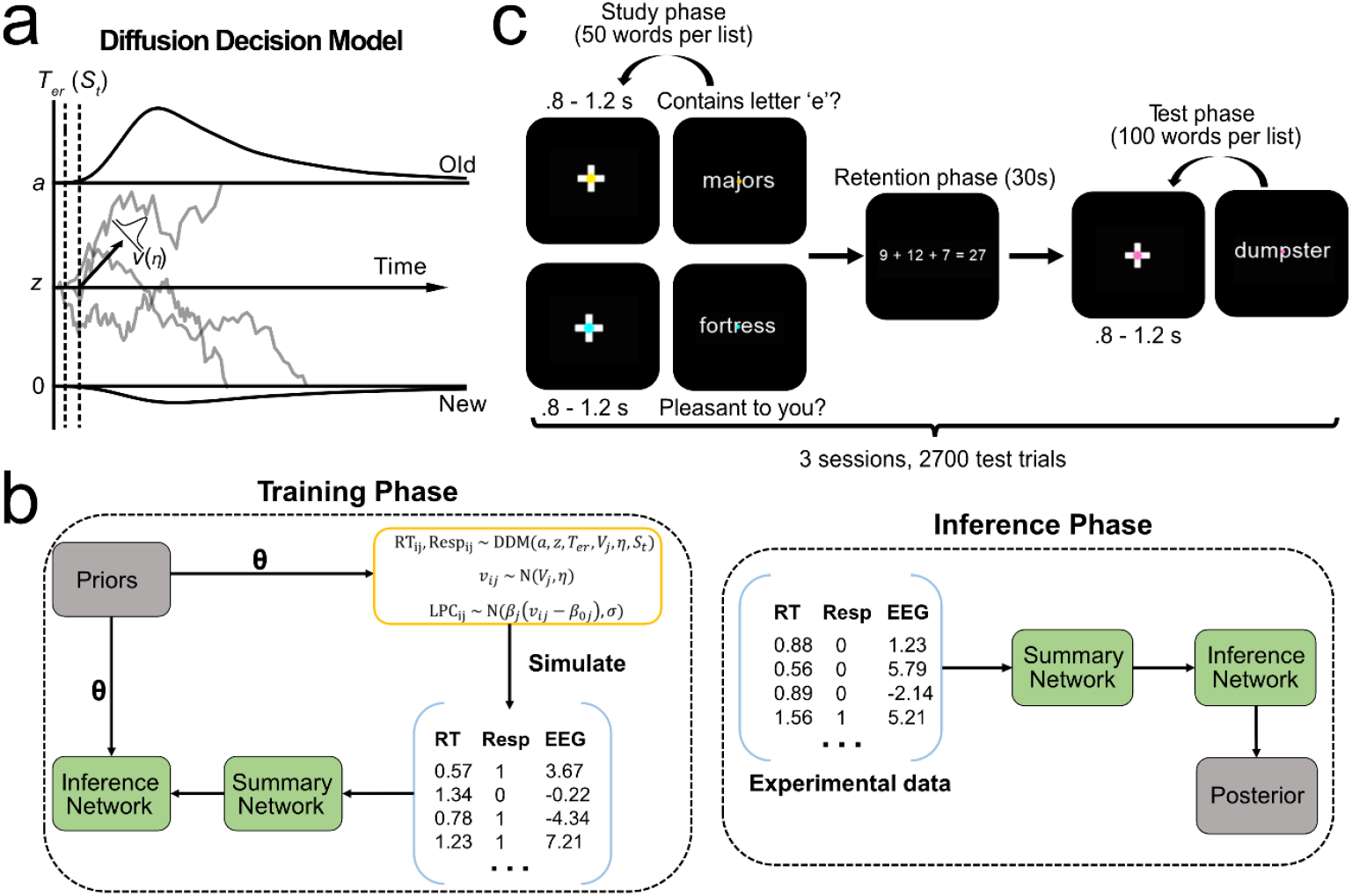
Diffusion Decision Model parameterisation, BayesFlow modelling pipeline and experimental structure. ***Note*. a.** Depiction of the Diffusion Decision Model for recognition memory tasks and conventional model parameters. Following word presentation, evidence accumulates in a stochastic manner toward the ‘old’ or ‘new’ decision boundary. *v* = drift rate, *a* = boundary separation, *z* = starting point, *η* = across-trial drift rate variability, *T*_er_ = non-decision time, *S*_t_ = non-decision time variability. **b**. An illustration of the BayesFlow workflow with a modified DDM for jointly predicting behavioural and neural data. Training was done simultaneously for the inference and summary networks by simulating the model with sets of parameters drawn from prior distributions. During the inference phase, the summary statistics from experimental data were passed to the inference network to produce posterior model parameter estimates. **c**. Trial diagram for the recognition memory task. Participants were presented with a list of words during the study phase. They were required to judge whether each word contained the letter ‘e’ (perceptual task) or whether the word was pleasant (semantic task). The task was indicated by the colour of the fixation stimulus. Participants then answered mathematical questions during a retention phase and then viewed a list of test items. They reported whether the item was ‘old’ or ‘new’ via speeded keypress.

Recent work has leveraged the high temporal resolution of EEG to study the neural correlates of evidence accumulation processes in the diffusion model (Brosnan et al., 2020; Kelly et al., 2021; Pisauro et al., 2017). In perceptual decision-making tasks, a key signature of evidence accumulation is the centro-parietal positivity (CPP), which exhibits a gradual, positive build-up that peaks around the time of response. The CPP has been proposed as a domain-general marker of the amount of accumulated evidence, independent of motor preparation (Kelly & O’Connell, 2013; O’connell et al., 2012b). The rate of CPP amplitude build-up has been formally linked to drift rates in the DDM (O’connell et al., 2012b; O’Connell & Kelly, 2021). Pre-response CPP amplitudes have also been shown to covary with decision accuracy (Feuerriegel et al., 2022), RT and confidence ratings (Grogan et al., 2023; Kelly et al., 2021; Ko et al., 2024; Philiastides et al., 2014) in circumstances where there may be trial-wise differences in the extent of evidence accumulation at the time of the decision.

Importantly, the LPC shows similar characteristics to the CPP, suggesting that it may reflect a similar evidence accumulation process in recognition memory tasks. Specifically, LPC amplitudes scale positively with decision accuracy and confidence (Addante et al., 2012; Rugg & Curran, 2007), and LPC amplitudes are also more positive-going for trials with faster RTs (van Vugt et al., 2019). Within the DDM framework, an increased likelihood of ‘old’ responses, faster RTs, and higher confidence ratings are predicted by a higher drift rate (i.e., a greater memory strength; Desender et al., 2021; Pleskac & Busemeyer, 2010).

In recent work, we demonstrated that the LPC is time-locked to the time of the response, like the CPP (Sun et al., 2024). Specifically, using the Residual Iteration Decomposition Toolbox (RIDE; Ouyang et al., 2015), we deconvolved the LPC into subcomponents time-locked to either the stimulus onset or the response. We observed that the typical left-parietal LPC ramped up and peaked relative to the time of the response, directly contrasting the widely held assumption that the LPC occurs around 400-800 ms after the probe onset. LPC amplitudes measured at left parietal electrodes also covaried with confidence ratings for both targets and lures. This suggests the LPC might represent a decision-making process during memory retrieval, such as mnemonic evidence accumulation in the DDM.

In this study, we investigated how features of the LPC relate to parameters in evidence accumulation models, and whether the functional correlates of the LPC can be reconceptualised within this modelling framework. Whilst earlier studies have typically correlated neural signals with DDM parameters at the group, individual, or trial level (Forstmann et al., 2008; Frank et al., 2015; Ratcliff et al., 2016; van Vugt et al., 2019), we used a joint modelling framework that simultaneously predicts behaviour and neural data from a shared decision process (Ghaderi-Kangavari et al., 2023). Specifically, LPC amplitudes were linked to trial-wise drift rates via a specified linking function, with explicit modelling of EEG-related noise—an aspect often overlooked in previous joint modelling implementations (Cassey et al., 2016; Turner et al., 2013). This allowed us to disentangle signal-driven variability in LPC amplitudes from noise and quantify how much across-trial LPC variation is attributable to drift-rate fluctuations. This method provides a mechanistic index of the cognitive variables driving LPC amplitude and a principled measure of relationship strength. If the LPC is a correlate of mnemonic-strength, we expect stronger associations between LPC amplitudes and drift rate for old than new items. If this relationship is selective, it should not generalise to other DDM parameters or to ERPs conceptually unrelated to memory strength. Moreover, if the LPC indexes a decision process that culminates immediately prior to a motor response, this relationship should be strongest in time windows nearest the response.

A critical technical limitation of fitting joint models using conventional techniques is the need to develop closed-form likelihood functions for each unique implementation. To address the potential limitation of intractability, we used a novel neural network method (BayesFlow) for amortised, likelihood-free inference (Radev et al., 2020). This method approximates the likelihood function via neural likelihood estimation, whereby specialised neural networks are trained to map summary statistics from model predictions to parameter values sampled from prior distributions (Fig. 1b). Beyond providing a solution to the intractability problem, this approach enables efficient posterior parameter estimation and has been successfully applied across disciplines (Radev, Graw, et al., 2021; Sokratous et al., 2023; von Krause et al., 2022; Wieschen et al., 2024).

We also assessed whether we would replicate our previous finding that the LPC is indeed time-locked to the response, and we examined how LPC amplitudes covary with memory-strength manipulations and RTs. In a recognition memory task (Fig. 1c), we manipulated both depth of processing (deep vs. shallow; Craik & Lockhart, 1972) and word frequency (Malmberg & Nelson, 2003). Both manipulations reliably influence recognition memory performance and LPC amplitudes; deep/semantic encoding results in higher successful identification rates and is associated with more positive-going LPC amplitudes than shallow encoding (Guo et al., 2004; Harris et al., 2013; Rugg et al., 1998). Low-frequency words are typically associated with higher hit rates, lower false alarm rates, and larger LPC old/new effects than high-frequency words (Bridger et al., 2014; Rugg, 1990; Rugg et al., 1995; Yang et al., 2019; Ye et al., 2019).

## 2. Method

### 2.1 Participants

We collected EEG data from 24 participants (19 female, 5 male, aged 18 – 33 years, M = 24.2, SD = 4). Participants were fluent in English and had normal or corrected-to-normal vision. One participant was left-handed. Two participants completed two out of the three sessions of data collection while four participants completed one session. No participants were excluded. All participants were paid 45 AUD per EEG session. This study was approved by the Human Research Ethics Committee of the Melbourne School of Psychological Sciences (ID 26845).

### 2.2 Stimuli

Stimuli were presented on a 24.5-inch ASUS ROG Swift PG258Q monitor (1920 x 1080 pixels, 60 Hz refresh rate). The stimuli were 900 words selected from UK-Subtlex database, which was an even split between low (2 - 5 per million) and high (80 - 500 per million) word frequency words. The word length was limited to 3 to 8 letters to reduce the need for making saccades. The word pool was re-used for each session but with random assignments to targets and lures. No words were repeated as stimuli within each session or list. There were 9 lists of words for the recognition task per session containing 50 targets and 50 lures, with balanced numbers of high and low frequency words. The stimuli were presented in lowercase and the same words were not repeated within each session as either targets or lures. Stimuli were presented using the Psychopy Toolbox (Peirce, 2007).

### 2.3 Paradigm

Participants were seated in a room with dimmed light and positioned 80 cm away from the monitor. For each session, participants completed a recognition memory task with EEG recordings. During the study phase, participants were instructed to view and memorise a list of 50 words presented serially at the centre of the screen. Each word was preceded by a fixation cross presented for 0.8 – 1.2 seconds (sampled from a uniform distribution). Each word was presented for 1.5 seconds along with a central fixation dot. Participants were instructed to fix their gaze on the fixation dot throughout the study phase.

Participants were additionally instructed to perform one of two tasks while memorising the word. They rated the pleasantness of the presented word (deep processing) for half of the study lists and indicated whether the presented word contained letter ‘e’ in it (shallow processing) for the other lists. The type of task to perform for each studied word was cued by the colour of the central fixation dot throughout the fixation and stimulus presentation periods (either blue or yellow, assignments counter-balanced across participants). Participants were instructed to complete the task by pressing ‘1’ or ‘2’ on a numpad using their index and middle fingers from one hand. The allocation of keys and responding hand were counter-balanced across participants.

Following the study phase, participants were instructed to complete a set of arithmetic problems for 30 seconds. An addition equation (e.g., ‘2 + 3 + 5 = 12’) was presented on the centre of the screen and participants indicated whether the equation was correct with a key response with the numpad (either ‘1’ or ‘2’). Feedback (i.e., ‘correct’ and ‘error’) were provided for each response for 1 s followed by a 0.5 to 1 second interval.

For the test phase, a total of 100 words were presented serially in the centre of the screen, containing 50 targets from the study list and 50 lures. Each word was preceded by a fixation cross for 0.8 – 1.2 seconds (sampled from a uniform distribution), Words were presented for a maximum of 5 seconds. Participants were instructed to indicate whether the presented word was old (from the last study list) or new using key responses with the numpad (either ‘1’ or ‘2’). If a response was made before the deadline, the word remained on screen for .5 seconds before the next fixation interval. Responses faster than .2 seconds or failures to respond within 5 seconds were immediately followed a screen displaying ‘too fast’ or ‘too slow’, respectively for 1 second. A pink fixation dot was presented in the centre of the screen throughout the test phase. As we used the same pair of keys (‘1’ and ‘2’) across the three tasks. For each participant, the key for indicating pleasant words or words containing ‘e’ during the study phase, ‘correct’ responses for the arithmetic task and ‘old’ responses for the recognition memory task were kept the same to avoid confusion.

A practice block was first completed using a list of 12 words for the study phase, and 24 words for the test phase. Feedback was provided (correct or wrong) for 1 s for each type of task in the practice block to help participants understand the instructions. Each practice list took about 4 minutes to complete, and participants had the choice to repeat the practice block if needed. Each experimental session included 9 blocks and 900 test trials in total. After EEG pre-processing, a total of 51822 trials (2159 trials on average per participant) were included in the final analysis.

To understand the performance in this task and test the effects of the experimental manipulations, we applied two-way repeated-measures ANOVA to test the main effects of word frequency and deep/shallow processing on hit rates for targets. For lures, we conducted t-tests to test the word frequency effects on correct rejection rates, given that the deep/shallow process distinction was not applicable to lures.

### 2.4 EEG data acquisition and processing

EEG data was recorded at a sampling rate of 512Hz using a 64-channel Biosemi Active Two system, using common mode sense (CMS) and driven right leg (DRL) electrodes (http://biosemi.com/faq/cms/&/drl.htm). We added six electrodes around the eyes, two placed 1 cm from the outer canthi of each eye, and four placed above and below each eye. We processed the data using EEGLAB (v 2021.1; Delorme & Makeig, 2004) running in MATLAB (The Mathworks). Code used for data processing and analysis will be available at https://osf.io/gqawp/ at the time of publication. First, we visually inspected the continuous EEG recording and excluded periods where EEG signals were excessively noisy. We also identified noisy channels (median number of noisy channels = 1, range = [0, 7]), which were excluded from the following data processing steps until they were interpolated. Second, we referenced the data to the average of all channels and low-pass filtered the data at 30 Hz (EEGLAB Basic Finite Impulse Response Filter New, zero-phase, -6 dB cutoff frequency 33.75 Hz, transition band width 7.5 Hz). We also removed channel AFz to account for the rank deficiency that can occur after the application of an average reference.

We applied Independent Component Analysis (ICA, RunICA extended algorithm) on a copy of the continuous EEG data (Jung et al., 2000) that was high-pass filtered at .1 Hz to improve stationarity of the signal. This allows for the reliable identification of ocular artifacts while minimising distortion of the blink components that can occur with stronger (e.g., 1 Hz) filters (Widmann et al., 2015). The resulting independent components were manually inspected, and components that resembled eye blinks and saccades were identified. These components were removed from the original continuous EEG data. Following this, we interpolated the noisy channels and channel AFz based on nearby channels (spherical spline interpolation). We then high-pass filtered the continuous EEG data at .1 Hz.

For the current study, we were only interested in EEG signals during memory retrieval. Therefore, the cleaned EEG data were segmented from -1,000 ms to 4,000 ms relative to the onset of each test probe. We then baseline-corrected the segmented data using a -200 to 0 ms time window prior to the onset of the test probe. As previous work (Sun et al., 2024) suggested that LPC should be best measured around the time of the response, we additionally constructed response-locked segments spanning -1,000 ms to 1,000 ms relative to the time of the response to each test probe, while using the same pre-stimulus baseline period. Please note that the response-locked data presented in the results section is after subtraction of the stimulus-locked subcomponent of the ERP, as described below.

### 2.5 ERP deconvolution and analyses

Following recommendations from previous work for measuring the LPC (Sun et al., 2024), we used the RIDE toolbox (Ouyang et al., 2011; Ouyang et al., 2015) to separate subcomponents of ERPs that are either time-locked to the onset of test probe or the response. Using the same parameter settings in previous work, we estimated two subcomponents: a stimulus-locked and a response-locked subcomponent, with time windows for estimation from 0 to 800 ms around stimulus onset, and -600 ms to 0 ms around response, respectively. For the input of the RIDE functions, we used the stimulus-locked epochs (-1,000 to 4,000 ms around stimulus onset) and estimated the subcomponent separately for each participant using pooled data across testing sessions. To reduce component overlap the stimulus-locked subcomponent was subtracted from all channels for analyses for response-locked data. This was done by subtracting the RIDE estimated stimulus-locked waveforms at each channel from each trial of EEG data before creating response-locked segments (Steinemann et al., 2018).

To identify the time windows that showed significant ERP amplitude differences, we performed mass univariate analyses with cluster-based permutation tests (Groppe et al., 2011; Maris & Oostenveld, 2007) implemented in the Decision Decoding Toolbox (Bode et al., 2019) to correct for multiple comparisons. This method groups adjacent time points with statistically significant amplitude differences between two conditions to form clusters of effects (cluster forming p < .01). The summed t values (the masses of the clusters) were then compared against the largest cluster masses obtained from each of 10,000 samples with permuted condition labels for a random subset of participants. The statistical significance of each cluster was judged by whether the cluster mass exceeded the 97.5th percentile of the permutation distribution (corresponding to an alpha level of 0.05, two tailed). For plotting purposes, we computed the within-participant standard errors across time points for comparing ERP waveforms (Morey, 2008).

To model changes in LPC amplitudes using the DDM, we calculated the averaged LPC amplitudes over a 200 ms time window prior to the time of the response. As the LPC effect has been consistently defined as a difference in EEG amplitude, we calculated the averaged amplitude over such a time window.

To visualise relationships between LPC amplitudes and RTs, we equally divided the EEG data within each participant into three equal tertiles based on the 33^th^ and 66^th^ percentiles, and then plotted the averaged waveforms from all participants based on both the stimulus-locked and response-locked data after subtracting the RIDE estimates. To investigate the association between pre-response LPC amplitudes and RT, we fitted a within-subject linear mixed effect model using RTs to predict LPC amplitudes in each trial, with random slopes and intercepts across participants. This was done for each sampling point within the 200 ms time window prior to the response for LPC. We then applied mass-univariate permutation tests to determine whether the *β* parameters obtained from linear mixed-effect models were significantly different to zero across time points.

### 2.6 Model parameterisation using BayesFlow

We employed BayesFlow to approximate likelihood functions for four versions of the DDM: a model linking LPC amplitudes with drift rate, a model predicting LPC amplitudes across three-time windows, a model predicting LPC and visual P1 amplitudes, and a model predicting LPC amplitudes based on non-decision time. Parameters shared by all three models are drift rate (*v*) varying according to stimulus type (old and new), deep/shallow processing and word frequency group, starting point *(z*), boundary separation *(a*), non-decision time (*T*_*er*_), the across-trial variability parameters for non-decision time (*S*_*t*_), and the drift rate variability parameter *(η*) varying across targets and lures. The four models differed in how the neural component was predicted (parameters in each model were detailed in Supplementary Table 1). The calculation of the pre-response LPC amplitude was based the on average over the time window where the LPC showed significant difference between hits and correct rejections. It is important to note that this choice should not contribute to a circular analysis (Kriegeskorte et al., 2009) in our modelling work. This is because we implemented separate sets of parameters for targets and lures that scaled drift rates to predict the LPC amplitudes, which independently predicts data related to hits and correct rejections. The general modelling pipeline was an extension of previous work proposing this method (Ghaderi-Kangavari et al., 2023) and details of each model are outlined below.

#### 2.6.1 Model 1: predicting LPC amplitudes based on drift rate

For the neural component of this model, LPC amplitudes were linearly predicted by model parameter with a scale (*β*) and an intercept (*β*_*0*_) separately for targets and lures, with a single measurement noise parameter *σ*. Parameter estimation from BayesFlow relies on the specialised neural network to learn the relationship between model parameters and data based on model simulations. Therefore, the model was simulated following the form below:

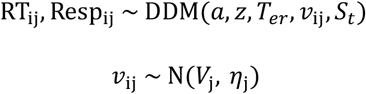

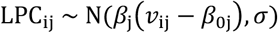

where j represents each experimental condition and i represents each trial. Therefore, this model assumed that choices and RTs were generated by the DDM with different drift rate parameters for each experimental condition. On each trial, the drift rate (*v*_ij_) was sampled from a normal distribution based on the mean drift rate estimated for each condition (*V*_j_) with a standard deviation of *η*. This drift rate was then linearly transformed as the mean of a normal distribution to generate the LPC amplitude on that trial, with a standard deviation of the noise parameter *σ*.

#### 2.6.2 Model 2: predicting LPC amplitudes in different time windows based on drift rate

For this model, we used drift rate sampled on each trial to simulate LPC amplitude across time windows via three separate functions with different scales 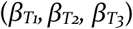 and intercepts parameters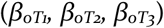. Further, this model included shared and independent noise across neural measures and data were simulated in a general form below:

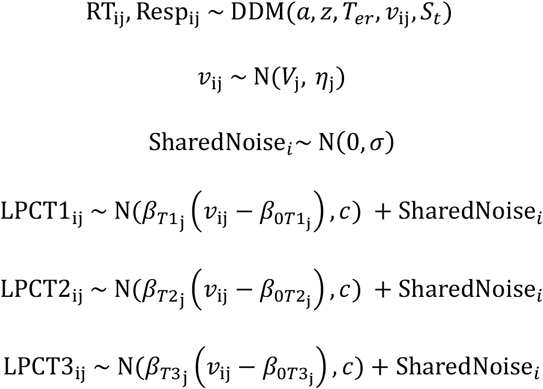

Whereas the shared noise component was the same for LPC in each time window signalling trial-level noise, it was assumed that LPC amplitudes in each 100-ms time window included unique noise that was not autocorrelated. This unique noise was simulated with an additional parameter *c*. Therefore, if the LPC amplitudes across three time windows were highly correlated, then we should expect a relatively smaller estimate of unique noise as compared to shared noise.

#### 2.6.3 Model 3: predicting LPC and P1 amplitudes based on drift rate

We measured the P1 as the mean amplitude from 50 to 150 ms relative to the stimulus onset at the electrode Oz. This model similarly assumed that LPC and P1 contains both shared and unique components of noise. Here, as LPC and P1 amplitudes were known to be associated with unique processes (memory vs. early visual processing), we included separate parameters for the unique noise component. EEG amplitudes for each measure were predicted with separate functions:

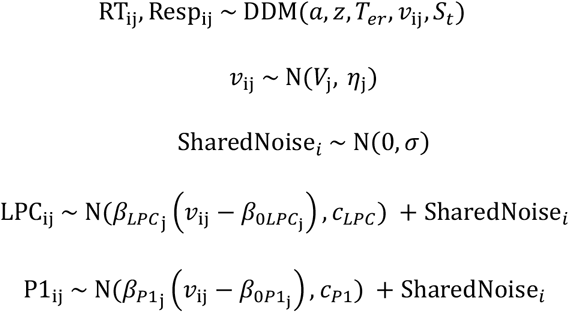

To investigate whether the specification of the sources of noise was sensible, we computed the amount of shared noise relative to the overall noise estimated in the data:

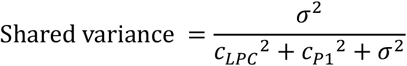

We then calculated the correlation between the amount of shared noise between the two EEG measures estimated by the model and the actual correlation across participants. It is expected that this measure should approximately track the trial-by-trial correlation between LPC and P1 amplitudes. This is because EEG is noisy, and it is expected that the neural/measurement noise should primarily contribute to the correlation between the two EEG component measures.

#### 2.6.4 Model 4: predicting LPC amplitudes based on non-decision time

This model was otherwise the same as the Model 1 except that LPC amplitude on each trial was predicted by the non-decision time parameter sampled from a uniform distribution.

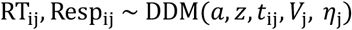

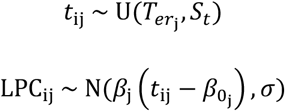

It is important to note that our simulation approach differed from studies that proposed the current BayesFlow method (Ghaderi-Kangavari et al., 2023), as we simulated data across conditions at the same time. In previous studies, the posterior parameters for different conditions were estimated by fitting a model without conditioned parameters multiple times to different parts of the data. This created multiple estimates of DDM parameters, such as boundary separation, that should be consistent throughout the experiment. This approach differs from traditional model fitting practices and provides less constraints on the parameter estimation. Further, we argue it is crucial in this case to have different parameters across conditions because numerous studies have shown that parameters such as drift rate and drift rate variability are different across targets and lures (Chen et al., 2024; Egan, 1958; Ratcliff et al., 1994).

For all models we trained the neural networks with experience-replay learning implemented in BayesFlow (Ghaderi-Kangavari et al., 2023). Experience-replay is technique widely applied in deep reinforcement learning that allows samples to be stored in a memory buffer and re-sampled for training (Zhang & Sutton, 2017), which has been suggested to improve data efficiency. The ranges of priors from which the parameters values were drawn are listed in the supplementary material. We trained each neural network with 1,000 epochs, 1,000 iterations per epoch, a batch size of 32 with a buffer size of 100. This means the network was trained on 32,000,000 partially repeating samples. For each simulated dataset, the number of trials varied from 300 to 500. While this number was smaller than the actual data size, we have demonstrated that such network can generate robust parameter estimation compared to the same model trained on a larger sample size (Supplementary Fig 8).

For parameter estimation, we generated 10,000 samples for each posterior distribution. For model predictions, we applied thinning and simulated data based on every 50^th^ sample of parameters in the posterior distribution (i.e., 200 data simulations in total). Importantly, we acknowledge that an inherent limitation of the neural network method is that the posterior estimates are informed by prior range but can sometimes fall outside of this range. To address this, we constrained the ranges for drift rate variability parameters and scale parameters with a logit-normal distribution. This involved first sampling unbounded parameter values from a normal distribution:

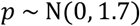

These parameter values were used by the network for learning. The choice of standard deviation of 1.7 was to make the logit-normal distribution to approximate a uniform distribution, consistent with the prior distribution assumption for other parameters. A demonstration of this transformation can be found in the supplementary materials (Supplementary Fig 9). Within the simulation functions, these parameters were transformed with a sigmoid function:

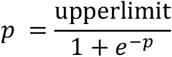

Therefore, for both the simulation and posterior estimation, the values of the parameters were constrained between zero and the pre-defined upper limit of the prior range, and sampled from a logit-normal distribution approximating a uniform distribution.

### 2.7 Calculating the Variance in EEG Measures Explained by Models

Following previous work (Ghaderi-Kangavari et al., 2023), we used parameters estimated from models to derive a variance-explained measure corresponding to the amount of variance in the EEG data explained by the model parameters. For example, for the model that linked LPC with single-trial drift rate, this ratio *r* was calculated as:

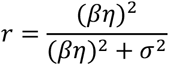

Where *η* is the across-trial drift rate variability parameter, *β* is the scale parameter of drift rate to produce the mean of LPC amplitude as a normal distribution with the noise parameter *σ*. For the model that linked LPC with non-decision time variability, the *η* parameter was replaced with the *S*_*t*_ for non-decision time variability. For the model that included additional independent source of neural noise, this measure was calculated based on the joint noise:

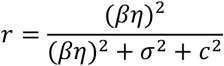

Where *c* represents another independent source of noise, such as the noise associated with P1 in Model 2. These estimates were produced based on parameter sets from the entire posterior distribution, the 95% highest density intervals of the measures were reported. We conducted statistical analyses (e.g., within-participant *t*-tests and repeated measures linear mixed effect models) based on the mean participant-level estimates to determine how they differ across conditions.

To clarify, we did not intend to interpret the variance-explained measure as the true signal-to-noise ratio of the LPC, given that EEG data was generated based on simplified distributional assumptions rather than models with plausible neural mechanisms. However, as demonstrated in the results section, this measure was highly useful to understand and compare associations between neural data and model parameters under the same modelling framework. Crucially, as demonstrated in the supplementary material, these parameters as well as the measures could be recovered by the model. Further, we found that the neural noise estimates in Model 3 could strongly track the across-trial correlation between different neural signals (Supplementary Fig.3c).

### 2.8 Model Parameter Recovery

To check whether the trained neural network can reliably generate posterior parameter estimates, we performed parameter recovery analyses. For each participant, we generated 200 simulations based on the median parameter estimates from the posterior distributions and re-fitted the model to each of the simulated datasets, generating 10,000 posterior samples. We used median estimates to avoid biases in the estimates by skewed posterior distributions, which was due to estimates of parameters being close to the prior boundary (e.g., low scale parameter estimates). We then plotted the median parameters against the 95% highest density interval for the recovered parameters. Specifically, given that we relied on the variance-explained measure for inferring associations between model parameters and neural measures, we also focused on examining the recovery of this measure.

## 3. Results

We recorded scalp EEG from 24 participants who each completed up to three sessions of a recognition memory task (Fig. 1c), with 900 test trials per session (maximum 2,700 trials per participant, 18 participants completed 3 sessions, 2 completed 2 sessions, 4 completed 1 session). Overall, participants performed well in the recognition task (mean hit rate = 64%, SD = 15%, mean correct rejection rate = 84%, SD = 13%). For hit rates, there was a significant main effect of deep/shallow processing, *F*(1, 23) = 23.41, *p* < .001, a significant effect of word frequency, *F*(1, 23) = 13.51, *p* = .001, but no significant interaction effect, *F*(1, 23) = 3.91, *p* = .06. Deeply processed items were associated with higher hit rates than shallowly processed items, mean difference = 16.5%. Compared to high frequency words, low frequency words had higher hit rates for targets, mean difference = 6.8%. For lures, low frequency words had higher correct rejection rates, mean difference = 5.0%, *t*(23) = 4.21, *p* < 0.001. Our results were in line with existing work using these manipulations (Craik & Lockhart, 1972; MacLeod & Kampe, 1996; Malmberg & Nelson, 2003).

### 3.1 The LPC is time-locked to the recognition memory decision

By comparing hits and correct rejections, we first replicated the conventional stimulus-locked old/new LPC effect (Fig. 2a; Rugg & Curran, 2007). The scalp distribution of the old/new effect was most prominent at left parietal electrodes. ERP amplitudes at channel P3 were more positive-going for hits between 533 and 895 ms after stimulus onset (Fig. 2a, critical cluster mass = 49.71, cluster mass = 668.35, *p* = .002). These findings were consistent with the conventional definition of the LPC (Rugg & Curran, 2007).

**Figure 2.**
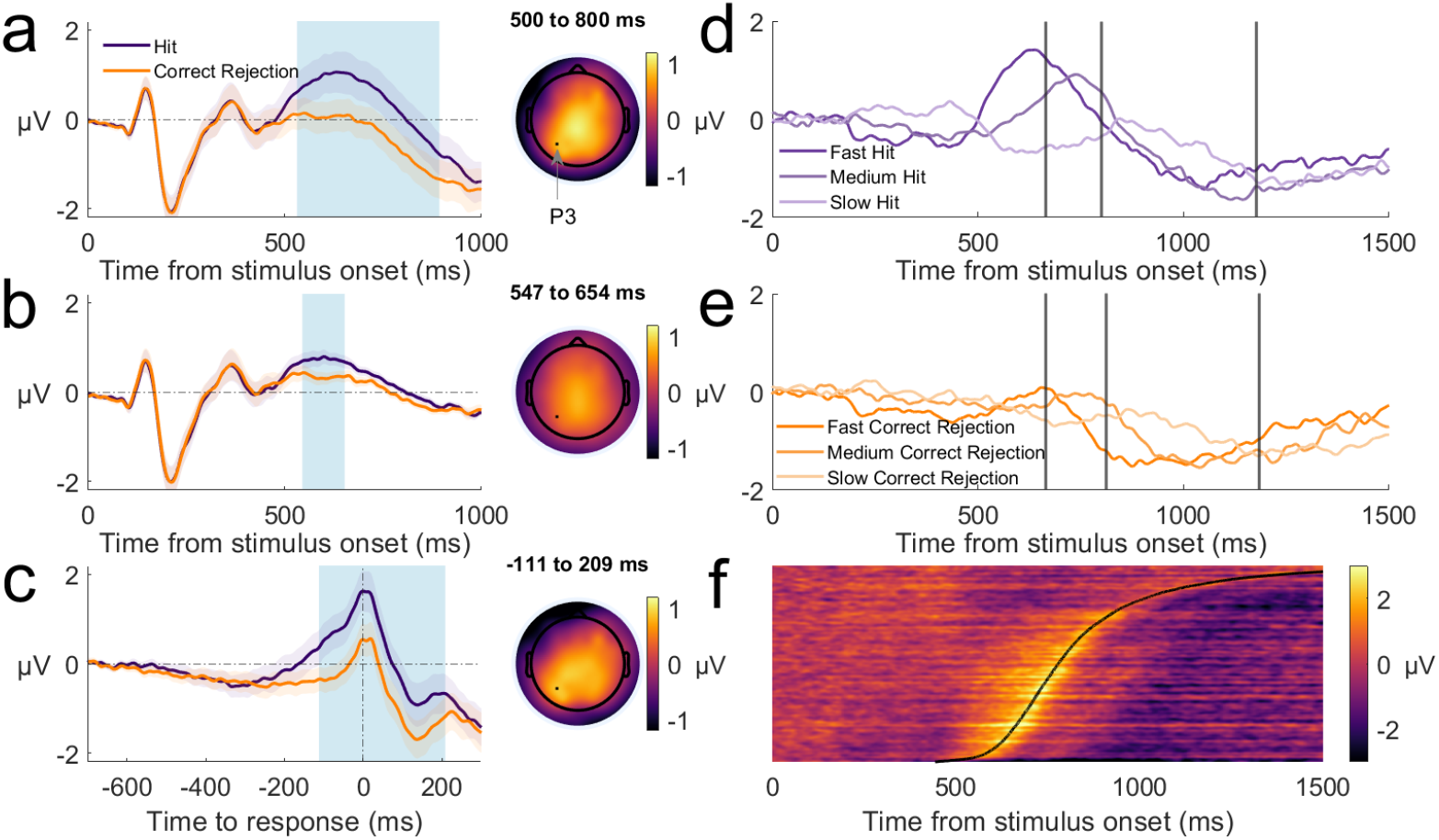
Stimulus- and response-locked ERPs in the recognition memory task. ***Note*. a.** The conventional LPC amplitude difference across hits and correct rejections (old/new effect), and the scalp distribution for hits minus correct rejections within the time window of statistically significant differences. Blue shading denotes time windows of statistically significant effects. **b**. The stimulus-locked ERP subcomponent estimated using RIDE and differences across hits and correct rejections. **c**. Response-locked LPC effects after subtracting stimulus-locked subcomponents from the data. **d**. Stimulus-locked ERPs for hits after subtraction of the stimulus-locked subcomponents, plotted by RT tertile bin. Vertical lines indicate average RTs for each bin. **e**. Stimulus-locked ERPs sorted by RT bin for correct rejections. **f**. ERPs for hits for all trials (pooled across participant) and sorted by RT, with the stimulus-locked subcomponents subtracted. Each row represents a single trial. Data were smoothed over 500-trial bins with a sliding Gaussian-weighted moving average filter.

A critical limitation of these stimulus-locked ERP analyses is that they also capture contributions from response-locked ERP components. Condition differences in the amplitude and/or timing of response-locked components can lead to spurious interpretations of stimulus-locked effects (Ouyang et al., 2017; Ouyang et al., 2015). Conversely, response-locked ERPs can also be biased by stimulus-locked ERPs (Frömer et al., 2024; O’Connell et al., 2025). To address this signal convolution problem, we separately estimated stimulus-locked and time-varying ERPs using the Residual Iteration Decomposition (RIDE) toolbox (Ouyang et al., 2015), as done in previous studies (Fong et al., 2025; Steinemann et al., 2018; Sun et al., 2024).

In the stimulus-locked ERP subcomponent, we observed a similar old/new effect between 543 and 659 ms post stimulus onset (Fig. 2b, critical cluster mass = 52.36, cluster mass = 202.53, *p* = .004). Consistent with our previous observations (Sun et al., 2024), this effect was small and demonstrated a central parietal origin, rather than the left-lateralised topography of the conventional LPC. The stimulus-locked subcomponent in each condition was subsequently subtracted from the single-trial ERPs (Steinemann et al., 2018). Response-locked segments of EEG data were then derived to better isolate the decision-locked signals.

In the response-locked data, the LPC gradually increased in amplitude and peaked at the time of the response (Fig. 2c). We observed more positive-going amplitudes for hits compared to correct rejections spanning -111 to 209 ms relative to the response (critical cluster mass = 64.93, cluster mass = 680.17, *p* < .001). This ramp-to-peak feature of the LPC was clearer for hits than correct rejections. We further demonstrate this feature in the stimulus-locked data with the stimulus-locked subcomponents subtracted. For both targets and lures (Fig. 2d and 2e), the timing of the LPC shifted further away from stimulus onset for slower RT tertiles as observed for the CPP (O’connell et al., 2012b). When sorting all trials in the experiment by RT (Fig 2f), the positive-going LPC signal is closely aligned with the time of the keypress response. Importantly, the topography of the response-locked LPC old/new effect showed a left-lateralised, parietal locus that is highly similar to the conventional stimulus-locked LPC effect (compare Fig 2a and 2c).

Taken together, this indicates that the LPC is time-locked to the response rather than the stimulus. It appears that the conventional stimulus-locked LPC effect arises from the contributions of overlapping, response-locked ERPs, where the RT distributions for hits and correct rejections closely align with the timing and duration of the conventional stimulus-locked effects.

### 3.2 Response-locked LPC amplitudes covary with accuracy and RT

Building on existing findings related to the conventional stimulus-locked LPC, we tested for associations between the response-locked LPC and behavioural performance (accuracy and RT). First, we found that response-locked LPC amplitudes were sensitive to our manipulation of memory strength. LPC amplitudes were larger for deeply compared to shallowly processed targets (Fig 3a, -157 ms to 173 ms to response, critical cluster mass = 36.06, cluster mass = 661.99, *p* = .001), consistent with the large behavioural effect of deep/shallow processing on hit rates. However, we did not observe a significant difference in the old/new effects between high and low frequency words, or a word frequency effect on the LPC amplitudes for hits and correct rejections (Fig. 3b and 3c). We expected that LPC amplitudes would be higher for less frequent targets and lower for less frequent lures, based on the observation that lower frequency words are more likely to be correctly recognised and less likely to be falsely recognised (Malmberg & Nelson, 2003). The non-significant effect could be due to the small behavioural effect observed for our word frequency manipulation (Fig. 4a).

**Figure 3.**
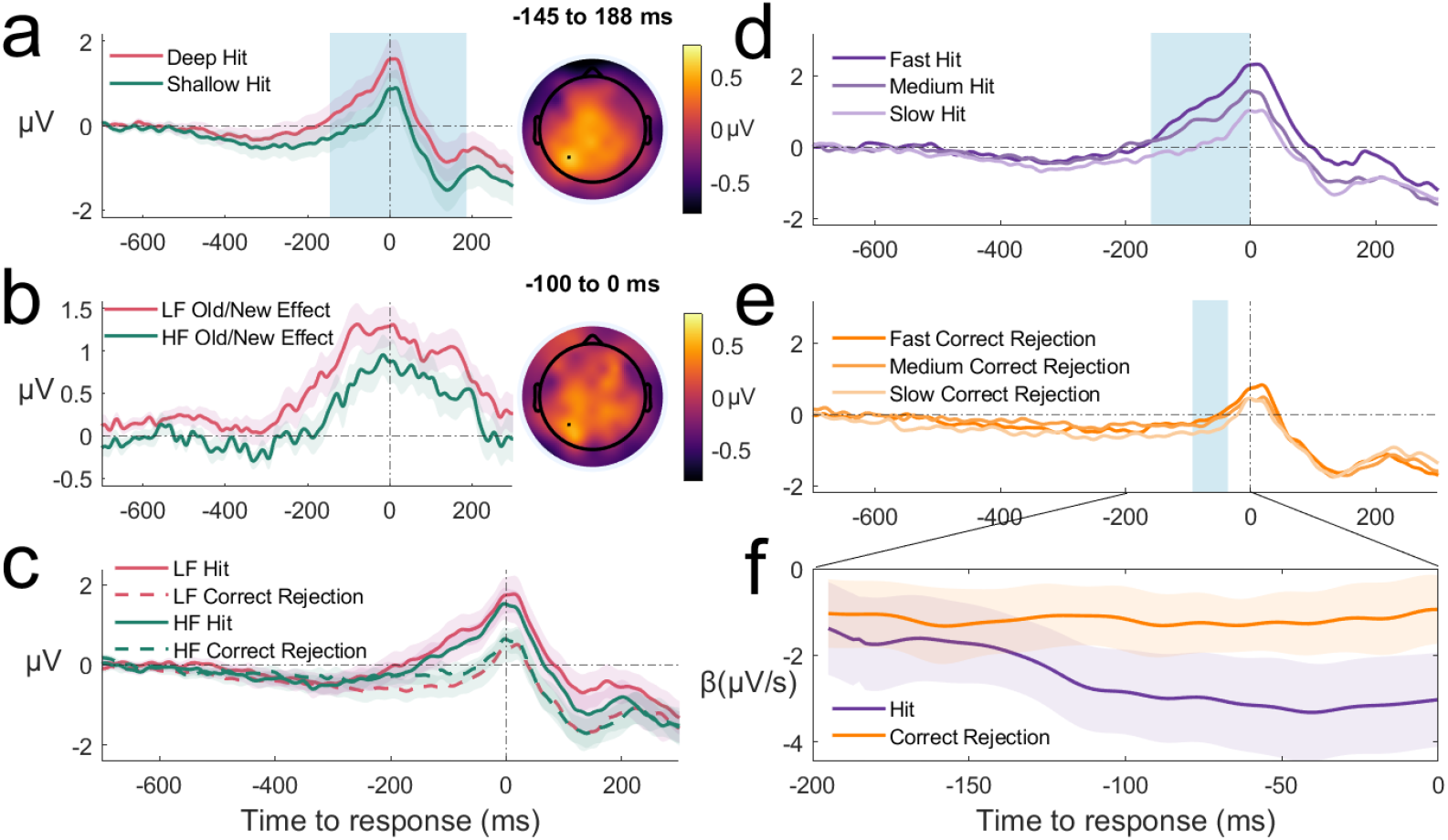
Response-locked ERPs by depth of processing, word frequency and RT. ***Note*.a.** Response-locked ERPs for hits in deep/shallow conditions, and the scalp distribution of deep minus shallow differences during the statistically significant time window marked by the blue shaded area. b. Response-locked ERPs for hits in high and low word frequency conditions, and the scalp distribution of the low minus high difference during 100-ms time window prior to the response. c. Response-locked ERPs for correct rejections in high and low word frequency conditions. d. Response-locked ERPs for hits binned by RT tertile [0-33%, 33-66%, 66-100%]. Blue-shaded areas denote statistically significant associations between RTs and LPC amplitudes. e. Response-locked ERPs for correct rejections by RT tertile. f. *β* parameters for the effect of RT on ERP amplitude estimated from linear mixed effect models. Shaded error regions depict within-participant standard errors for each plot. LF = Low Frequency. HF = High Frequency

**Figure 4.**
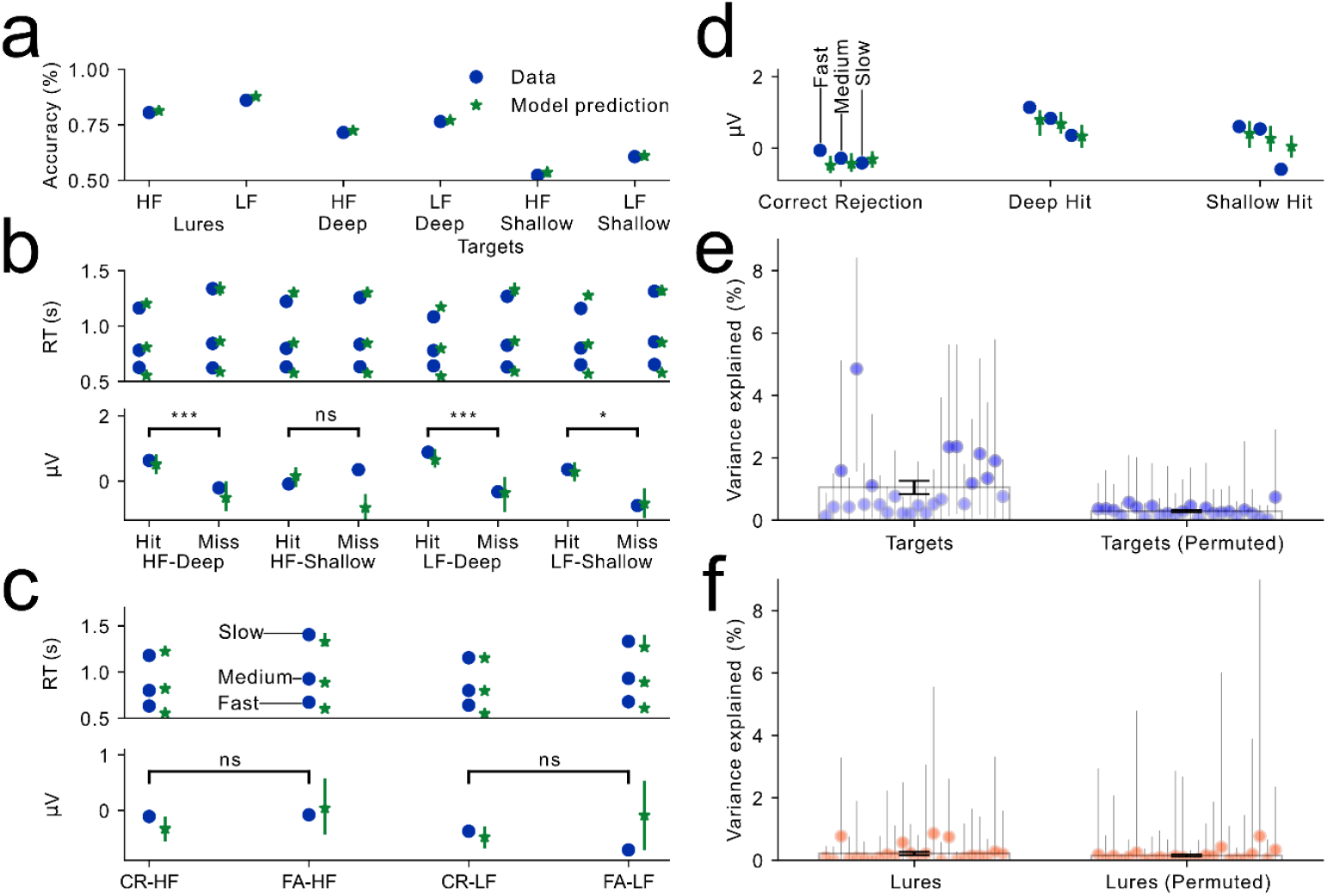
Model predictions and estimated variance-explained measures from the model using drift rates to predict LPC amplitudes. ***Note*. a.** Model 1 predictions of accuracy across conditions. The blue dots represent data averaged across participants. The green stars represent model predictions averaged across participants over 200 posterior samples (every 50^th^ sample in the posterior distribution). Error bars denote 95% highest density intervals. b. Model 1 predictions of RT percentiles and mean LPC amplitudes for targets across word frequency, processing depth and hit/correct rejection responses. c. Model 1 predictions of RT percentiles [10^th^, 50^th^ and 90^th^ percentiles] and mean LPC amplitudes for lures across high/low word frequency conditions and correct rejection/false alarm responses. d. Model 1 predictions for the LPC amplitudes averaged within equally binned RT tertiles [0-33%, 33-66%, 66-100%]. e. LPC amplitude variance-explained measures for targets and permuted data. Dots represent the median estimates for participants. Error bars depict 95% highest density intervals estimated from 10,000 posterior samples. f. Variance-explained measures for lures. ns = non-significant, * = *p* < 0.05, *** = *p* < 0.001. LF = Low Frequency. HF = High Frequency. CR = Correct Rejection. FA = False Alarm

We also found that response-locked LPC amplitudes increased with faster RTs for both targets and lures (Fig. 3d and 3e). This partially explains the shifts and decreases in the LPC amplitude from fast to slow RTs in the stimulus-locked data (Fig. 2d and 2e). The *β* parameters from mixed effects models across time points were significantly lower than zero across time points prior to the response, indicating lower LPC amplitudes for faster RTs. Associations were stronger for hits than correct rejections, as evidenced by visual inspection and *β* values (Fig. 3f). Notably, for hits, the association was stronger at time points closer to the response.

Overall, LPC amplitudes were larger in conditions associated with a higher memory strength and for faster responses, and this effect was larger for targets. These characteristics are consistent with changes in the drift rate across conditions and trials within the DDM, where a higher drift rate will more likely result in faster RTs and a higher likelihood of ‘old’ responses.

### 3.3 DDM predictions account for patterns of behavioural and neural data

To better understand the relationships between the LPC and behaviour, we jointly modelled accuracy, RT and response-locked LPC amplitudes using a modified DDM (Model 1). Specifically, LPC amplitude on each trial was sampled from a normal distribution, where the standard deviation of the distribution reflects the noise in the neural measure and the mean is linked to the drift rate from the DDM via a linear function (see Fig 1B). As a more positive-going LPC amplitude has been associated with more accurate recognition (Rugg & Curran, 2007; Rugg & Yonelinas, 2003; Yonelinas, 2002), we assume that LPC amplitudes positively scaled with drift rate. To construct a single measure of LPC amplitude for each trial, we calculated the averaged amplitude in a 200-ms time window prior to the response. We used BayesFlow (Version 1.0; Radev et al., 2020) for amortised inference and parameter estimation.

We then fitted the model to the experimental data and generated 10,000 samples for each posterior distribution of parameters. The DDM was able to capture patterns of accuracy scores and RT distributions (summarized by the 10^th^, 50^th^, and 90^th^ percentiles) across conditions in the experiment (Fig 4a-4c). The unique advantage of our joint modelling framework is its ability to capture relationships between accuracy/RT and LPC amplitude – in addition to capturing mean LPC amplitude across conditions that differ in accuracy (Fig 4b and 4c). Specifically, the model predictions closely match the mean LPC amplitudes across RT tertiles (Fig 4d), which showed a stronger relationship for targets than lures. We also found that the median parameters for posterior distributions across participants could be well recovered (Supplementary Fig. 1a). Notably, we also replicated the typical finding of a greater across-trial drift rate variability for targets than lures (Chen et al., 2024; Starns & Ratcliff, 2014). This demonstrates the advanced ability of the neural network to approximate likelihood functions for complex models.

Visual inspections of the model predictions against data (Fig 4b and 4c) indicated two main places of misfits for the neural data: LPC amplitudes between correct rejections and false alarms, and hits and misses for shallowly processed high word frequency targets. Here, we note that these apparent numerical differences did not correspond to statistically significant differences in our ERP measures. Specifically, there were no significant differences in LPC amplitudes between correct rejections and false alarms for either high or low frequency lures (*p* = 0.837 and *p* = 0.373), or between hits and misses for shallowly processed high word frequency targets (*p* = 0.379). Therefore, our model was able to capture changes in the neural data that corresponded to statistically significant ERP effects.

### 3.4 LPC amplitudes covary with single-trial drift rates

We next assessed whether LPC amplitudes covaried with drift rate estimates in each trial. An advantage of our modelling approach is the consideration of noise in EEG data, which could be understood as variance in EEG that was not explained by the specified model parameters. This parameterisation additionally allowed the estimation of the relative contributions to the LPC variance estimates from across-trial drift rate variability and noise. We used the median variance-explained values from the posterior distribution for statistical analysis. Across participants, we found this median variance-explained measures were significantly higher for targets than lures within participants (Fig 4e and 4f), *t*(23) = 4.30, *p* < 0.001. To validate this link between LPC and drift rate across trials, we permuted the trial-level match between LPC amplitudes and behavioural data across participants. Variance-explained measures were significantly lower than those from unpermuted data for targets, *t*(23)= 3.58, *p* = 0.002, but not for lures, *t*(23)= 1.16, *p* = 0.260, with the latter estimates close to zero. Therefore, the LPC amplitudes covaried with drift rate estimates in our models, and this association was much stronger for targets than lures.

While the variance-explained measures might be considered low (1.07 % on average for targets), within the context of the amount of noise in the data, they are substantial. This was demonstrated by estimating the maximum explainable variance in the LPC measures based on split-half consistency (Bainbridge, 2017; Hebart et al., 2020; Tarhan et al., 2021). We first randomly split the participants in two halves and calculated the mean LPC measures across repeated test probes for each half. We then calculated the correlation of the measures across the two halves, for either target or lure words, and applied Spearman-Brown correction to the coefficients. These coefficients were estimates of maximum explainable variance (or “noise ceiling”). With 10,000 iterations of randomly selected halves, we found that averaged estimates were 4.22% (95% HDI [1.22%, 8.44%]) for targets and 5.05% (95% HDI [1.92%, 8.93%]) for lures. While these group-level measurements do not necessarily apply for each participant, this suggested that only a small amount of variance in the LPC was explainable, given the highly noisy EEG recordings. Despite the same method can be used to calculate the noise ceiling for each participant, it is not feasible in the current dataset given the low numbers of repeated trials for each participant. The relatively high level of noise in single-trial EEG signals is also demonstrated by the small size of the LPC old/new effect (mean = 1.01 µV) relative to standard deviation of LPC measures across trials (9.08 µV). Nevertheless, the estimates observed here were of similar magnitude to the same estimates observed in a previous recognition memory experiment (Ghaderi-Kangavari et al., 2023). These measures could also be well recovered by our model (Supplementary Fig 1b).

### 3.5 The relationship between the LPC and drift rate is strongest around the time of the response

To further understand how associations between LPC amplitudes and drift rates changes over time, we modified the DDM to simultaneously account for LPC amplitudes from three 100-ms time windows of LPC leading up to the time of the response (Model 2). The LPC amplitude in each time window was independently predicted by the drift rates across trials. To account for the autocorrelation in EEG data, the three LPC time windows shared trial-level noise but also contained independent noise across time windows. Using a linear mixed-effect model we found that the later time windows (-200 to -100 ms, -100 to 0 ms) showed significantly greater variance-explained estimates than the -300 to -200 ms time window, *β* = 0.67, *p* = 0.02, *β* = 2.20, *p* < 0.001. A significant main effect of stimulus type was also observed, with the estimates being greater for targets than lures, *β* = 0.62, *p* = 0.007. This indicated a stronger association between LPC amplitudes and drift rates over time towards the time of the response across targets and lures. Notably, trial-level noise was estimated to be more dominant than across-window noise (Supplementary Fig 2a), consistent with the autocorrelation assumption.

Notably, we observed that the LPC slope, estimated from -300 to 0 ms around the response, provided a higher variance-explained measure than the LPC amplitude (Supplementary Fig 11). The same slope measure has been used as a key feature of CPP. However, we showed that this association could be partially accounted by the LPC amplitude at the response.

### 3.6 ERPs unrelated to mnemonic strength do not account for drift rate variance

We tested the selectivity of the relationship between the drift rate and the LPC by modifying our joint model to also include measures of the visual P1 component (Model 3), measured at around from 50 to 150 ms after stimulus onset at electrode Oz. This component is generally assumed to reflect early visual processing across different tasks (Di Russo et al., 2002; Smith et al., 2003). We therefore did not expect P1 to be strongly associated with memory strength. For the model specification, LPC and P1 amplitudes were predicted by drift rate in one model independently, and the noise was split into shared and independent components. Overall, we found that the LPC explained a decent amount of the variance in drift rate, but not the P1.

Specifically, using a linear mixed effect model we found a significant main effect of stimulus type, with targets associated with significantly higher variance-explained measures than lures, *β* = 1.60, *p* = 0.001. We also found a significant interaction between ERP component and stimulus type, with the difference between P1 and LPC being larger for targets than lures, *β* = -1.95, *p* = 0.004. For lures, the variance-explained measures did not differ significantly between LPC and P1, *β* = 0.03, *p* = 0.945.

As a validation to Model 3, we observed that the independent components of the EEG noise were more dominant than the shared noise (Supplementary Fig 3a). This was consistent with the fact that the time windows between the LPC and P1 components were further apart than the LPC measures in the three-time-window model. Further, across participants, we found the estimated proportion of shared noise over overall noise showed a near-perfect correlation (*r* = 0.96, *p* < 0.001) with the correlation coefficients estimated between the LPC and P1 across trials (Supplementary Fig 3c). This suggests that the current assumption of Gaussian noises was able to track across-trial correlation of the outcome variables, which provides strong evidence that inferences from the model were based on information across trials.

We also tested the spatial selectivity of the response-locked ERP with drift rate. Specifically, under the same modelling framework, we let the model to jointly predict the pre-response LPC amplitude and another concurrent measure at the frontal channel F6. The choice of the electrode was to reduce overlapping with another previously reported FN400 component at the central frontal regions (Rugg & Curran, 2007). We found that the LPC showed stronger association with drift rate than the response-locked measure at F6 (Supplementary Fig 10), providing additional support for the selectivity of the relationship.

### 3.7 The LPC is not related to variability in non-decision time

Finally, we validated our findings by testing whether changes in LPC amplitudes could be explained by other parameters in the DDM (Model 4). We chose the non-decision time parameter (Ratcliff et al., 2004; Ratcliff & Tuerlinckx, 2002), which similarly varies across trials (see Fig. 1). This model was otherwise the same as the first model except that LPC amplitude on each trial was predicted by the non-decision parameter sampled from a uniform distribution. Results from fitting such a model showed that non-decision time across trials could not account for model predictions of LPC amplitudes (Supplementary Fig 7) and the median variance-explained measures were close to zero (Fig. 5c). All three models mentioned so far could well account for the neural data (except for the non-decision time model) and behavioural data, and the parameters could be well recovered (Supplementary Figs 2 to 7).

**Figure 5.**
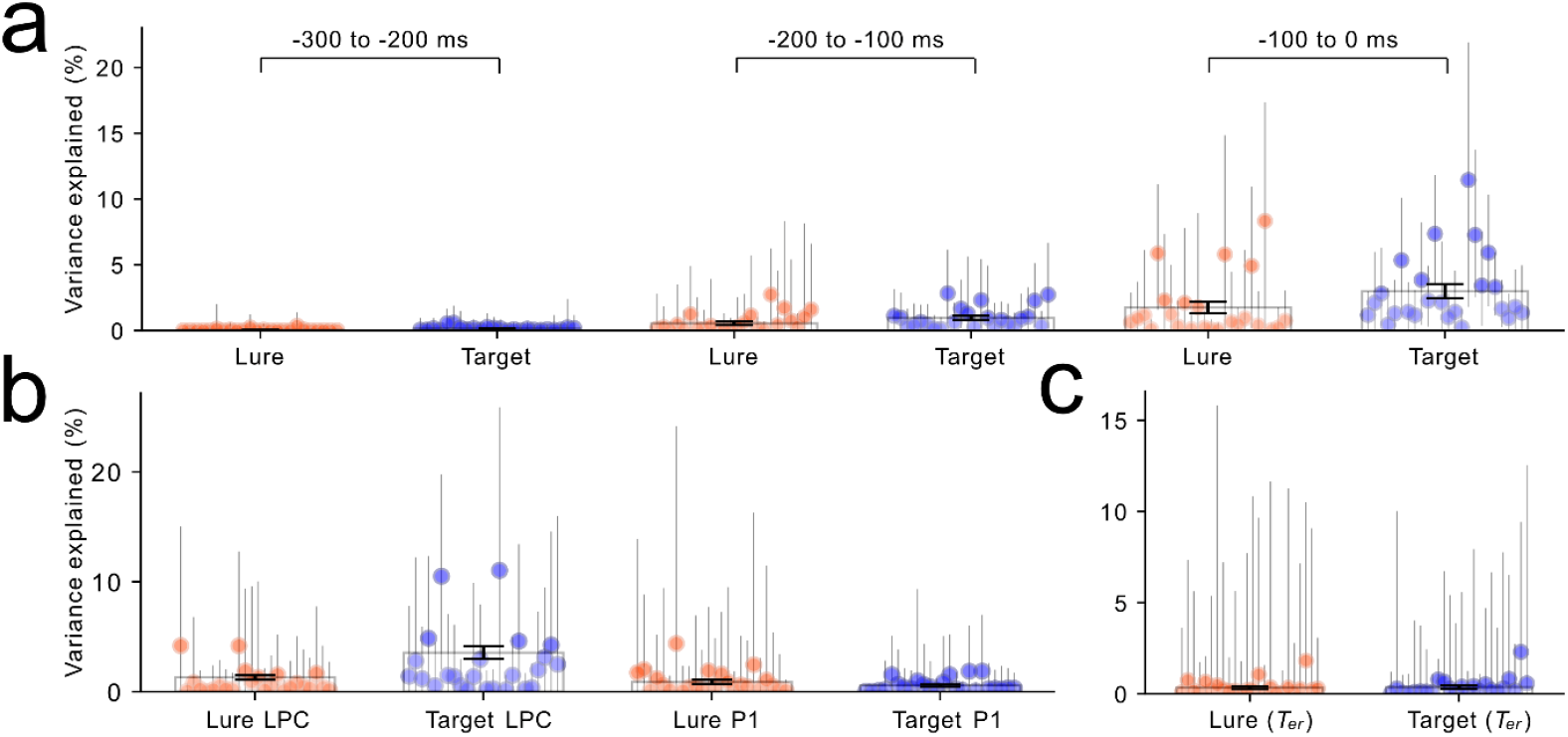
Variance explained measures from the validation models. ***Note*. a.** LPC amplitude variance explained measures estimated for lures and targets across three time windows leading up to the response based on Model 2. b. Variance explained measures estimated for lures and targets for LPC and visual P1 component amplitudes based on Model 3. c. Variance explained measures estimated for lures and targets for the LPC based on non-decision time variability based on Model 4.

## 4. Discussion

Inspired by the mnemonic accumulator hypothesis, we used a joint modelling framework to investigate whether the parietal LPC is a neural correlate of evidence accumulation rate in the DDM. We built on our recent findings (Sun et al., 2024) by demonstrating that the LPC is more accurately characterised as a response-rather than a stimulus-locked ERP component. We also observed properties of the LPC that are consistent with a mnemonic strength signal. Using generative modelling to jointly account for behaviour and EEG signals, we found that pre-response LPC amplitudes were associated with across-trial variation in drift rate, indexing mnemonic strength, and we found that the LPC was selectively associated with variations in drift rate, but was not with a non-mnemonic factor that varies across trials (non-decision time).

Our findings offer a fundamental reinterpretation of the LPC as a neural correlate of item memory strength driving evidence accumulation during recognition memory decisions. This contrasts with the conventional assumption that the LPC represents a high-threshold recollection process during memory retrieval (Kwon et al., 2023; Rugg & Curran, 2007; Yonelinas, 2002), with LPC amplitudes indexing either successful or failed recollection. Diminished LPC amplitudes were often interpreted as reflecting impairment in retrieving memory details (Kim et al., 2004; Olichney et al., 2008; Wolk et al., 2013). By directly integrating LPC measures into a computational model that specifies an evidence accumulation process, we show that the LPC instead reflects a continuously varying mnemonic strength signal, with larger amplitudes associated with faster decisions, contrary to the assumption of recollection as a slow process (Rugg & Curran, 2007). While other theories proposed that recollection could also be operationalised as a continuous variable (Onyper et al., 2010; Wixted & Mickes, 2010), it is important to note such proposals have largely overlooked response times.

This alternative interpretation of the LPC is consistent with the mnemonic accumulator hypothesis (Wagner et al., 2005), which proposes that the parietal cortex integrates mnemonic evidence rather than representing memory details, and particularly asymmetrically accumulating evidence for ‘old’ but not ‘new’ decisions (Sestieri et al., 2017; Sestieri et al., 2014). The asymmetry hypothesis was based on observations that parietal BOLD signals and single-neuron recordings selectively track accuracy and confidence for old responses (Henson et al., 1999; Hutchinson et al., 2015; Kahn et al., 2004; Rutishauser et al., 2018; Wheeler & Buckner, 2003) and lesion studies showing selective impairments for “old” but not “new” decisions (Hower et al., 2014). In our data, the LPC showed a clearer ramp-to-peak waveform and stronger pre-response associations with RT and drift rate across models for targets than for lures. Therefore, a more positive-going LPC amplitude could be associated with greater memory strength and faster accumulation.

While our modelling results provided new evidence for this asymmetry, we suggest further investigation is warranted due to other previous ERP findings and plausible accounts of the current ERP observations. First, other studies have also reported findings consistent with a symmetrical accumulator in the parietal lobe, showing increased parietal activity as measured via single-neuron recordings and BOLD signals, for both high-confidence “old” and “new” decisions (Rutishauser et al., 2018; Sestieri et al., 2014). Likewise, more positive-going parietal ERP amplitudes are sometimes observed for high (as compared to low) confidence “old” and “new” responses (Brezis et al., 2017; Rubin et al., 1999; Woodruff et al., 2006; Woroch & Gonsalves, 2010). We have also observed similar effects in response-locked data (Sun et al., 2024) with a midline centro-parietal component that was different to the left-lateralised LPC. Second, while we observed a lack of association between the LPC and RT for lures, this could be due to the lower evidence variability for lures (see Chen et al., 2024; and current variability estimates in Suppmentary Table 2-5). Further work is needed to determine the extent to which current observations reflect genuine asymmetry in mnemonic evidence accumulation. This will require disentangling different sources of parietal neural activities and implement stronger manipulations of memory variability for lures.

### 4.1 Are the LPC and the CPP equivalent to each other?

Given the established link between the LPC and mnemonic evidence accumulation, it seems reasonable to ask – is the LPC in recognition memory isomorphic to the CPP, the evidence accumulation signal found in perceptual decision-making tasks? The two share some resemblance in that they both have morphologies that co-vary with decision accuracy (Feuerriegel et al., 2022), RT, and confidence (Grogan et al., 2023; Sun et al., 2024). Here we list several salient differences in the ERP waveforms that prevent us from endorsing a view that they reflect the same underlying signal.

First, in contrast to the LPC, the CPP shows a slower build-up of amplitude that onsets earlier relative to the motor response in trials with slower as compared to faster RTs (Feuerriegel et al., 2021; Fong et al., 2025; Kelly & O’Connell, 2013). This corresponds to a longer averaged evidence accumulation trajectory with slower build-up for trials with lower drift rates. Second, the CPP converges to a relatively fixed amplitude prior to the response (Kelly & O’Connell, 2013; O’connell et al., 2012a), which is consistent with a fixed decision threshold. In our analysis, we instead observed that LPC amplitudes around the time of the response were more positive-going for faster RTs in hit trials, and were associated with drift rate. We did not observe an earlier onset of positive-going ERP amplitude build-up for trials with slower RTs. Third, while the CPP is typically observed to be similar in amplitude across choice options in perceptual two-choice tasks, in our data, we observed an asymmetry across old and new responses. It should be noted that the LPC and CPP are measured not only across different tasks but also several different task parameters including different stimulus biases and difficulties. Future work should also attempt to equate these task parameters before concluding that the two ERP components reflect different cognitive aspects of decision-making.

So what does the LPC reflect exactly? On one hand, it shows a clear relationship to drift rate in that larger LPC amplitudes are observed with faster RTs around the time of the response. On the other hand, the LPC is response-locked and shows its strongest relationship to drift rate at times closest to the response. The latter two points are somewhat odd when one considers that in the conventional DDM, the drift rate is fixed throughout the duration of the trial. Thus, if the LPC reflects a process that *determines* the drift rate, it should show its strongest association to drift rate after the stimulus encoding time, but it should not show a stronger relationship after that point.

Before we speculate, we would like to clarify that our results most clearly indicate that the LPC is a *correlate* of drift rate rather than a direct measure of it. With this in mind, we can propose some possibilities that could be explored in future work. The first possibility is that our LPC measurements index a process that closely covaries with RT and drift rate, which is not necessarily a process that determines the drift rate. For example, previous work has identified ERP correlates of decision confidence that are prominent at the time of the motor response (Fukuda et al., 2024; Sun et al., 2024). Similar correlates of confidence have also been observed in perceptual tasks (Grogan et al., 2023; Ko et al., 2024). These effects exhibit a midline-focused scalp amplitude distribution that differs from the typically left-lateralised LPC but would nevertheless influence LPC measurements at left parietal electrodes via volume conduction. Neuroimaging methods with better source localisation, such as magnetoencephalography informed by structural magnetic resonance imaging data, may be able to better disentangle contributions from different cortical sources.

The second possibility is that the memory strength being indexed by the LPC is not constant as the standard DDM assumes, but increases throughout the course of the trial. In some DDM variants, the drift rate is not fixed throughout the duration of the trial, but instead begins at a low value and increases to an asymptotic value as the trial proceeds (Smith & Lilburn, 2020; Smith & Ratcliff, 2009). While these models are commonly applied to perceptual decision-making, an analogous class of recognition memory models assumes that the cues used to make contact with memory are gradually assembled throughout the course of the trial, leading to stronger evidence around the time of the response due to more complete cues (Brockdorff & Lamberts, 2000; Cox & Shiffrin, 2017). According to these models, our results may instead suggest that memory strength itself is increasing throughout the trial.

It should be mentioned that these two possible explanations are merely speculation, as our current data and models cannot easily adjudicate between them. While it would be possible to apply time-varying drift rate models to our data, such models would likely require additional manipulations to characterize the drift rate trajectories, such as time pressure manipulations (Smith & Lilburn, 2020) or a response-signal procedure (Brockdorff & Lamberts, 2000).

### 4.2 Future directions

While we demonstrated LPC fluctuations to reflect trial-level mnemonic strength, our framework does not use characteristics of the target or lure stimuli to understand the sources of single-trial drift rates. In recognition memory, a global matching process was proposed where the cue item is compared with memory representations of all studied items. The sum of all the pairwise similarity (i.e., the global similarity) represents the memory strength of the cue item. This could be an example of the integration processes that gradually compute memory strength. Recent development of such models derive item memory strength from stimulus similarity (e.g, semantic and orthographic similarities of words, Chang et al., 2025; Osth & Zhang, 2024; Reid et al., 2026; Zhang & Osth, 2024), which could be correlated with LPC amplitudes to inform model parameters. Although such approaches show promise— particularly for false recognition (Osth & Dennis, 2020; Osth, Zhang, & Williams, 2025; Reid & Jamieson, 2023)—current models using external measures of similarity have yet to provide a good account of the variability in memory strength for correctly recognised targets (see Nosofsky et al., 2025). Using these models, future studies could further inform the computation of memory strength with neural measures of the internal states of the decision-maker (Nunez et al., 2017; Sun et al., 2025).

Our study also demonstrates the benefits of advanced methods for integrating neuroimaging and computational modelling. We applied EEG deconvolution (RIDE) to better isolate response-locked ERP waveforms (Steinemann et al., 2018; Sun et al., 2024), mitigating concerns about temporal overlap with stimulus-locked signals (Frömer et al., 2024; O’Connell et al., 2025). We also leveraged a neural network-based, likelihood-free inference method (Ghaderi-Kangavari et al., 2023) to jointly model behaviour and EEG signals, enabling robust hypothesis testing within a unified framework. Specifically, we extended this method to simultaneously account for data across experimental manipulations, and predict multiple neural observations. The application and validation of these complex models further demonstrated the capability of network-based likelihood-free inference tools. As these tools continue to develop (Fengler et al., 2021; Lenzi et al., 2023; Sainsbury-Dale et al., 2024; Zammit-Mangion et al., 2024) and evolve—enabling functions such as model comparison (Radev, D’Alessandro, et al., 2021) and misspecification detection (Schmitt et al., 2023)—they hold promise for more precise mechanistic insights into cognitive processes and their neural correlates.

## 5. Conclusion

Our study provides compelling evidence that the LPC, traditionally seen as a marker of recollection, reflects a dynamic decision variable linked to mnemonic evidence accumulation during recognition judgments. We showed that LPC amplitudes are selectively associated with across-trial variations in drift rate within the DDM, consistent with the asymmetrical mnemonic accumulator hypothesis.

Methodologically, we demonstrate the advantages of using deconvolution methods to disentangle stimulus and response-locked signals, and a joint modelling framework to simultaneously fit behavioural and neural data. Further work in this field could further benefit from refining item-level inference and leveraging multimodal data to further validate the functional correlates of EEG waveforms in recognition memory tasks.

## 6. Data and Code Availability

Data and code used for data processing and analysis will be available at https://osf.io/gqawp/ at the time of publication.

## 7. Author Contributions

Conceptualization: JS, DF, AFO

Methodology: JS, DF, MN, AFO

Investigation: JS, DF, MN, AFO

Visualization: JS

Supervision: DF, AFO

Writing—original draft: JS

Writing—review & editing: DF, MN, AFO

## 10. Acknowledgements

We would like to thank Dr. Michael Nunez for sharing their code and for their guidance relating to model fitting using BayesFlow.

## 10. Supplementary Material

**Supplementary Figure 1.**
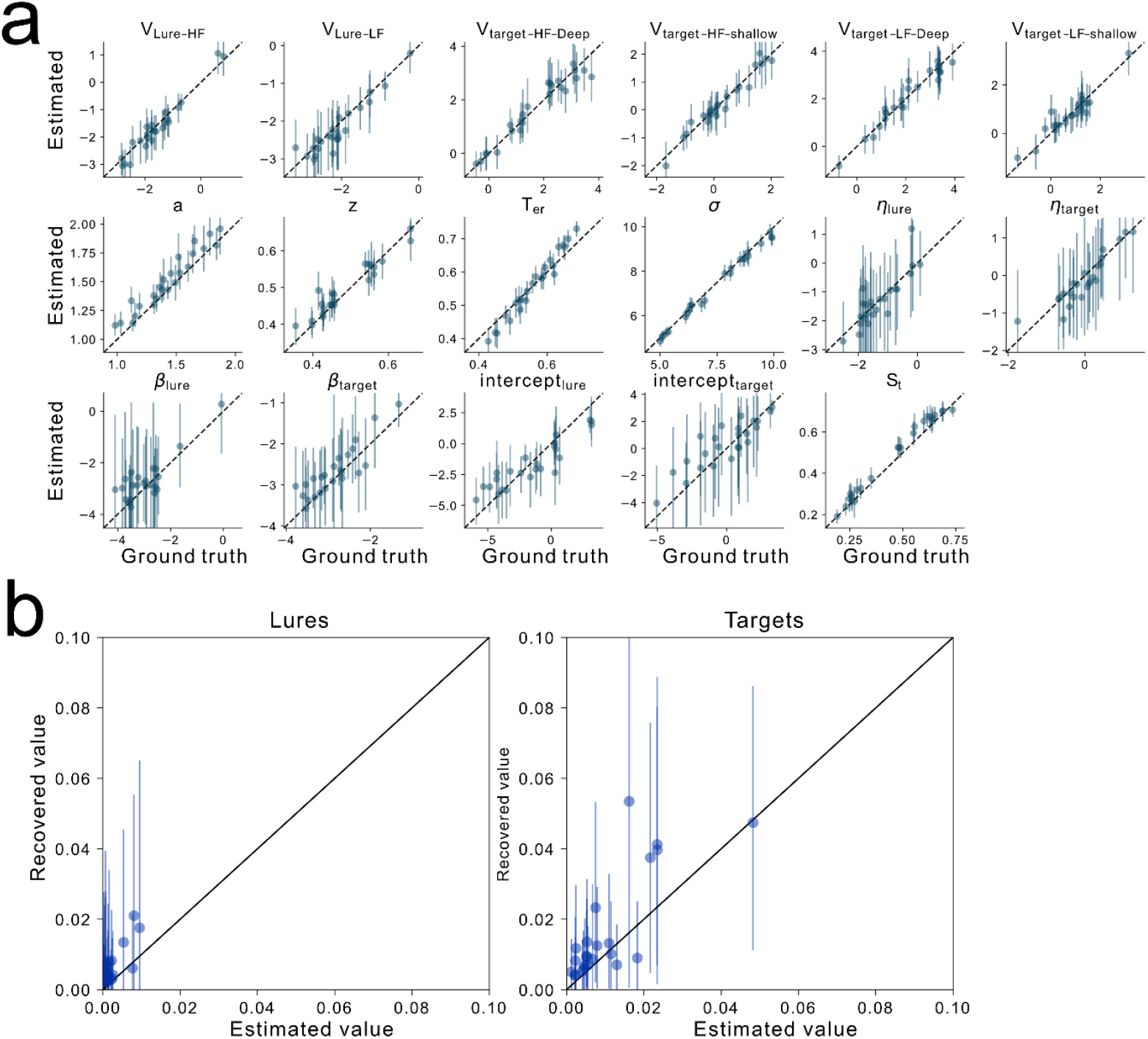
Recovery of best fitting parameters and variance-explained measures from the model that links LPC amplitude with drift rate. ***Note***. a. Parameter recovery of the best fitting parameters for each participant. The best fitting parameters were the most frequent in the posterior distributions, which were plotted against the mean of the recovered parameters across 500 samples. The error band represents the 95% highest density interval for the recovered means for parameters across the 500 samples. b. Recovery of the variance-explained measured across 500 samples.

**Supplementary Figure 2.**
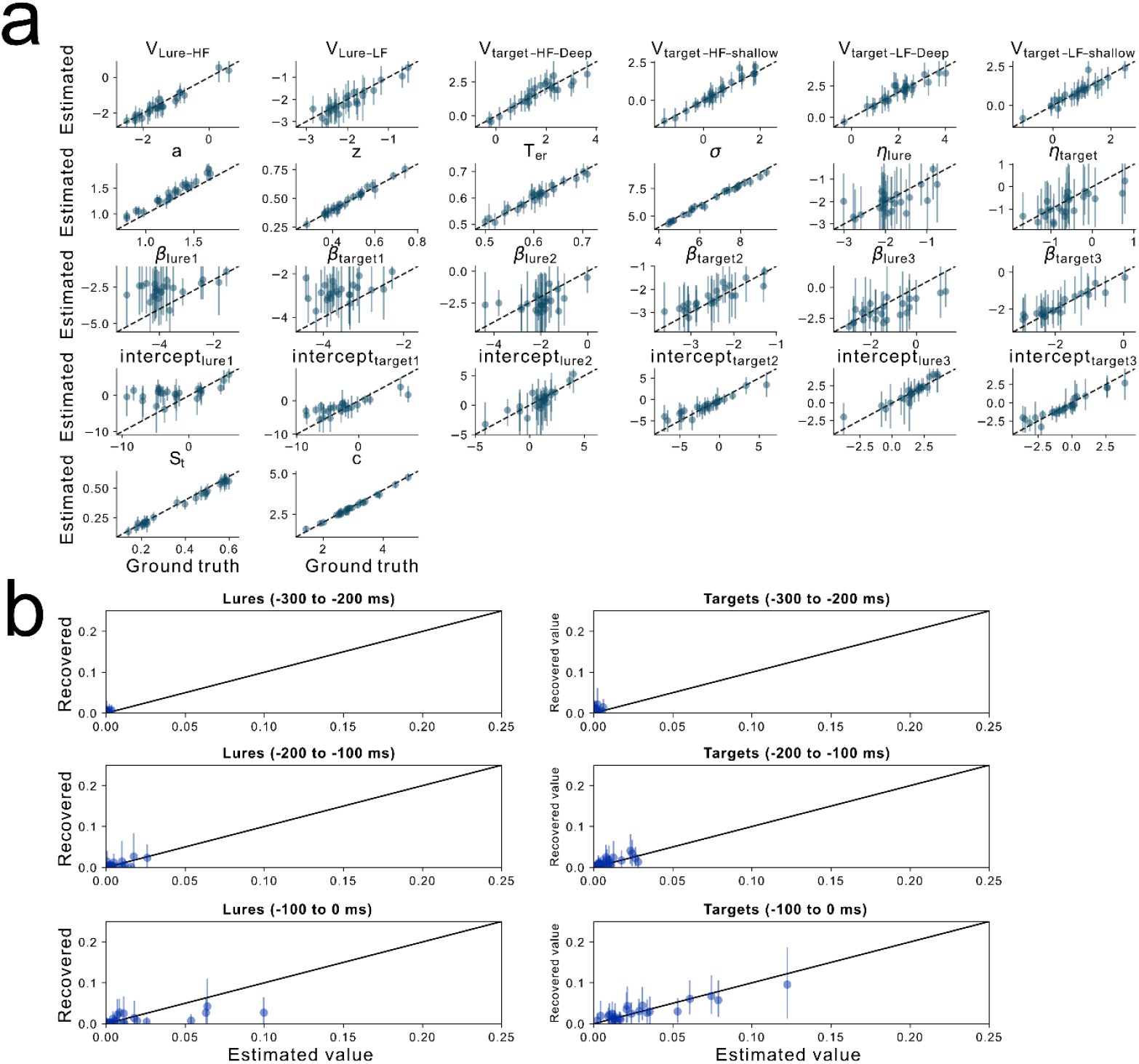
Recovery of best fitting parameters and variance-explained measures from the model that links LPC amplitudes in three time windows with drift rate. ***Note***. a. Parameter recovery of the best fitting parameters for each participant. The best fitting parameter were the most frequent in the posterior distributions, which were plotted against the mean of the recovered parameters across 500 samples. The error band represents the 95% highest density interval for the recovered means for parameters across the 500 samples. b. Recovery of the variance-explained measured across 500 samples for each time window.

**Supplementary Figure 3.**
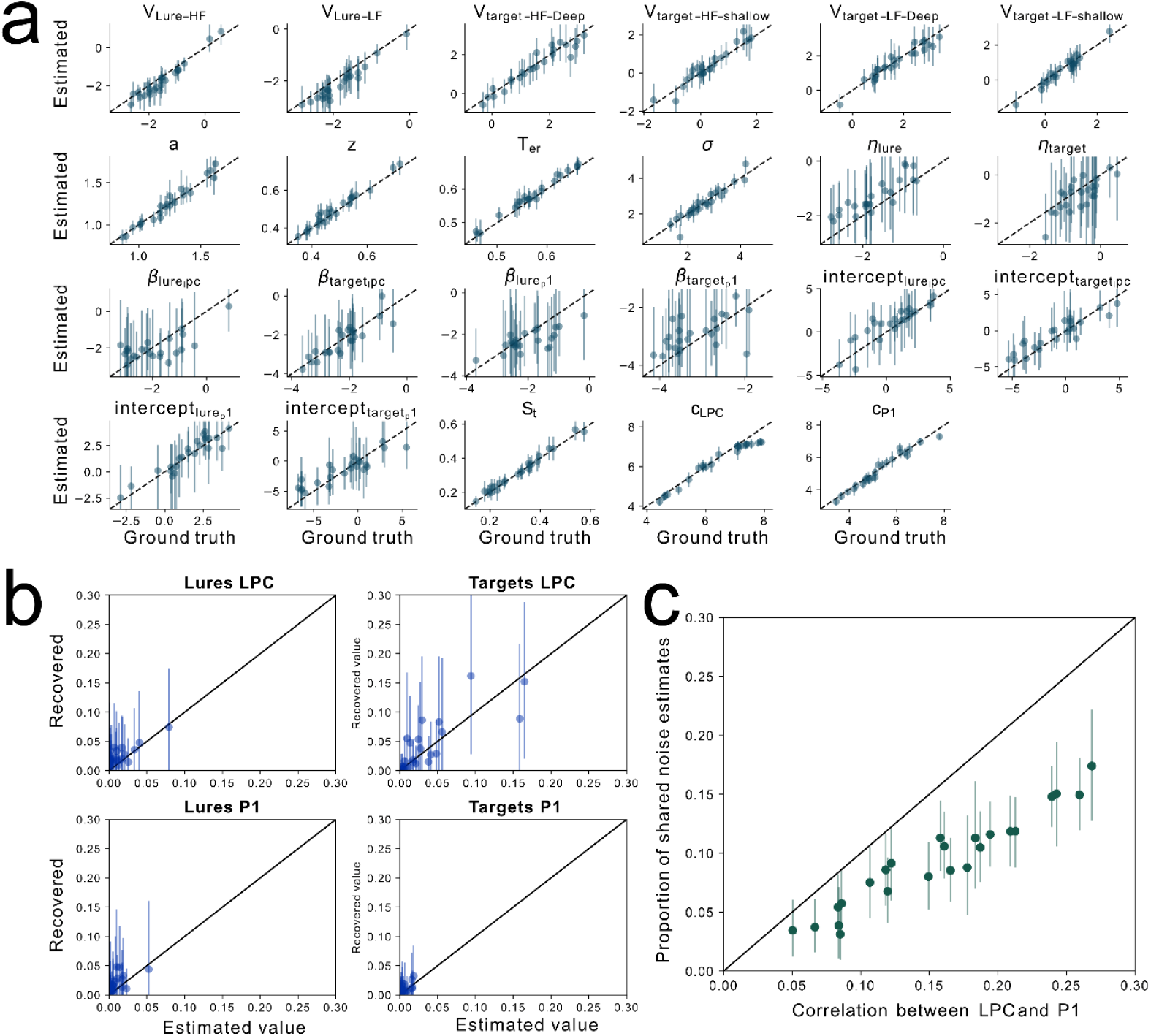
Recovery of best fitting parameters and variance-explained measures from the model that links drift rate with LPC and visual P1 amplitudes. ***Note***. a. Parameter recovery of the best fitting parameters for each participant. The best fitting parameter were the most frequent in the posterior distributions, which were plotted against the mean of the recovered parameters across 500 samples. The error band represents the 95% highest density interval for the recovered means for parameters across the 500 samples. b. Recovery of the variance-explained measured across 500 samples for LPC and P1visual P1 amplitudes. c. The correlation between the model estimated shared noise between LPC and P1 and the trial-by-trial correlation between LPC and P1 across participants.

**Supplementary Figure 4.**
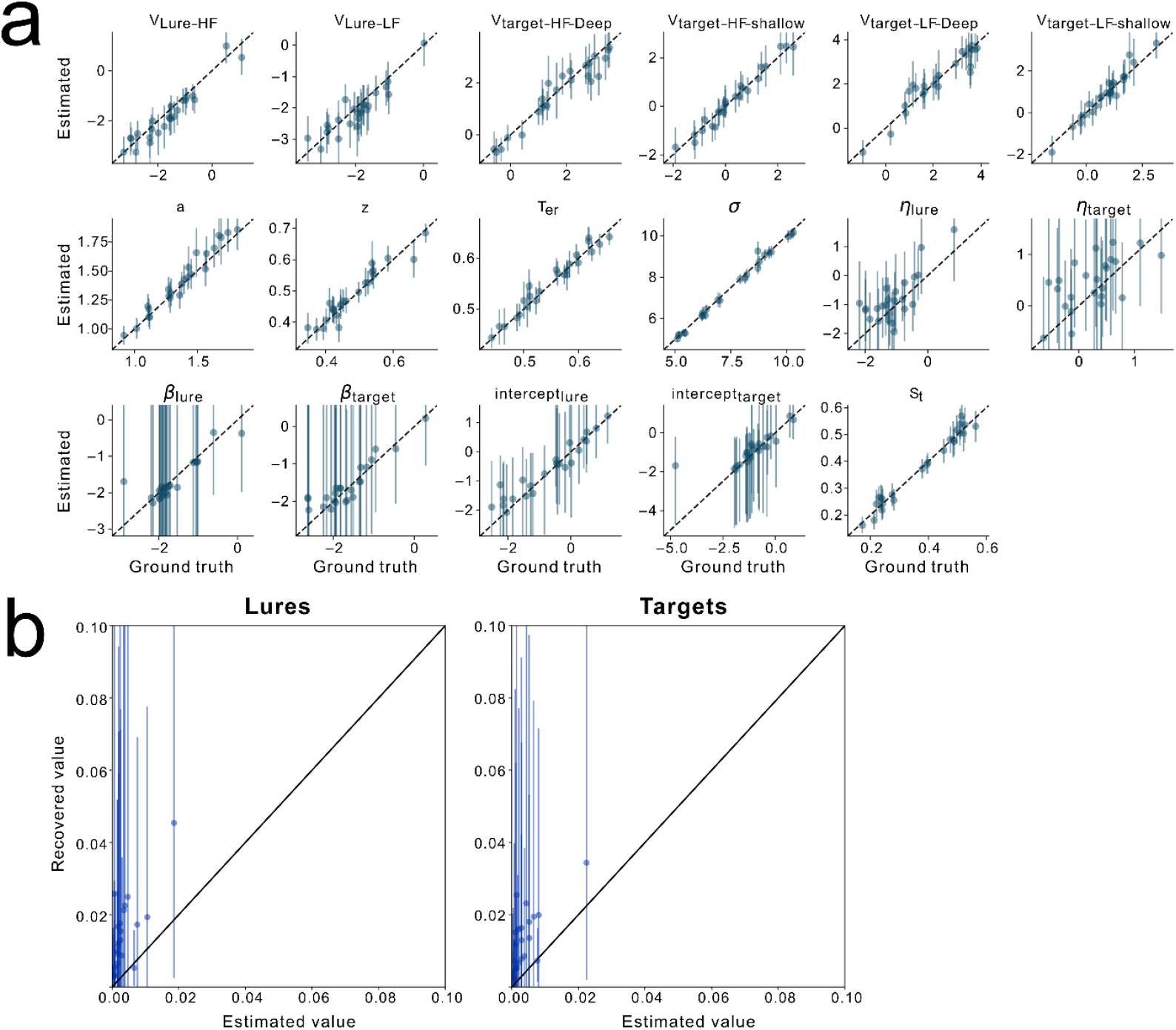
Recovery of best fitting parameters and variance-explained measures from the model that links non-decision time with LPC amplitudes. ***Note***. a. Parameter recovery of the best fitting parameters for each participant. The best fitting parameter were the most frequent in the posterior distributions, which were plotted against the mean of the recovered parameters across 500 samples. The error band represents the 95% highest density interval for the recovered means for parameters across the 500 samples. b. Recovery of the variance-explained measured across 500 samples.

**Supplementary Figure 5.**
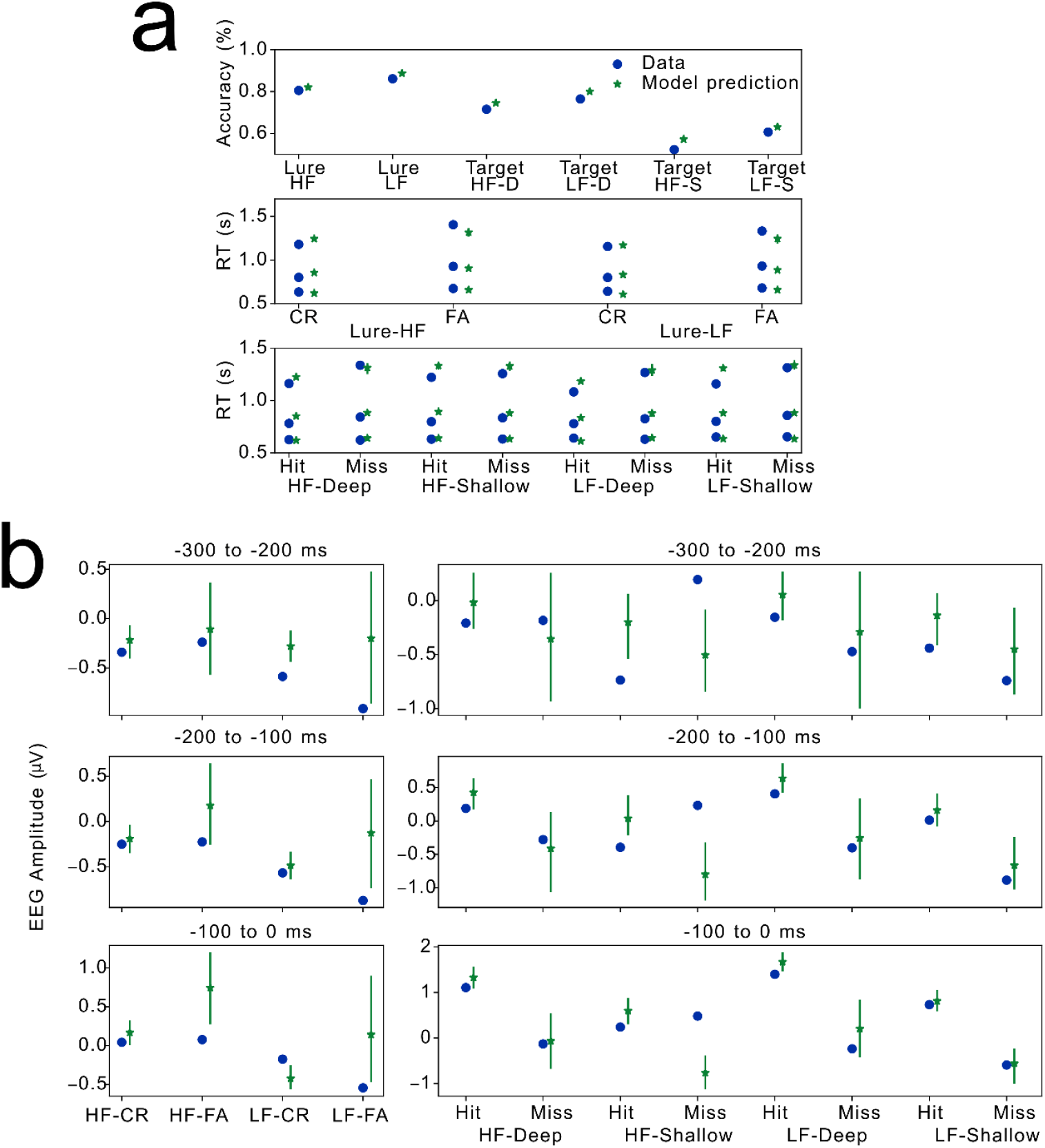
Model predictions for the model linking drift rate with three time windows of LPC amplitudes. ***Note***. a. Model predictions of behavioural data based on the best fitting parameters across participants. b. Model predictions of mean LPC amplitude across three time windows data based on the best fitting parameters across participants. The error band represents the 95% highest density interval across 200 samples generated from the best fitting parameters.

**Supplementary Figure 6.**
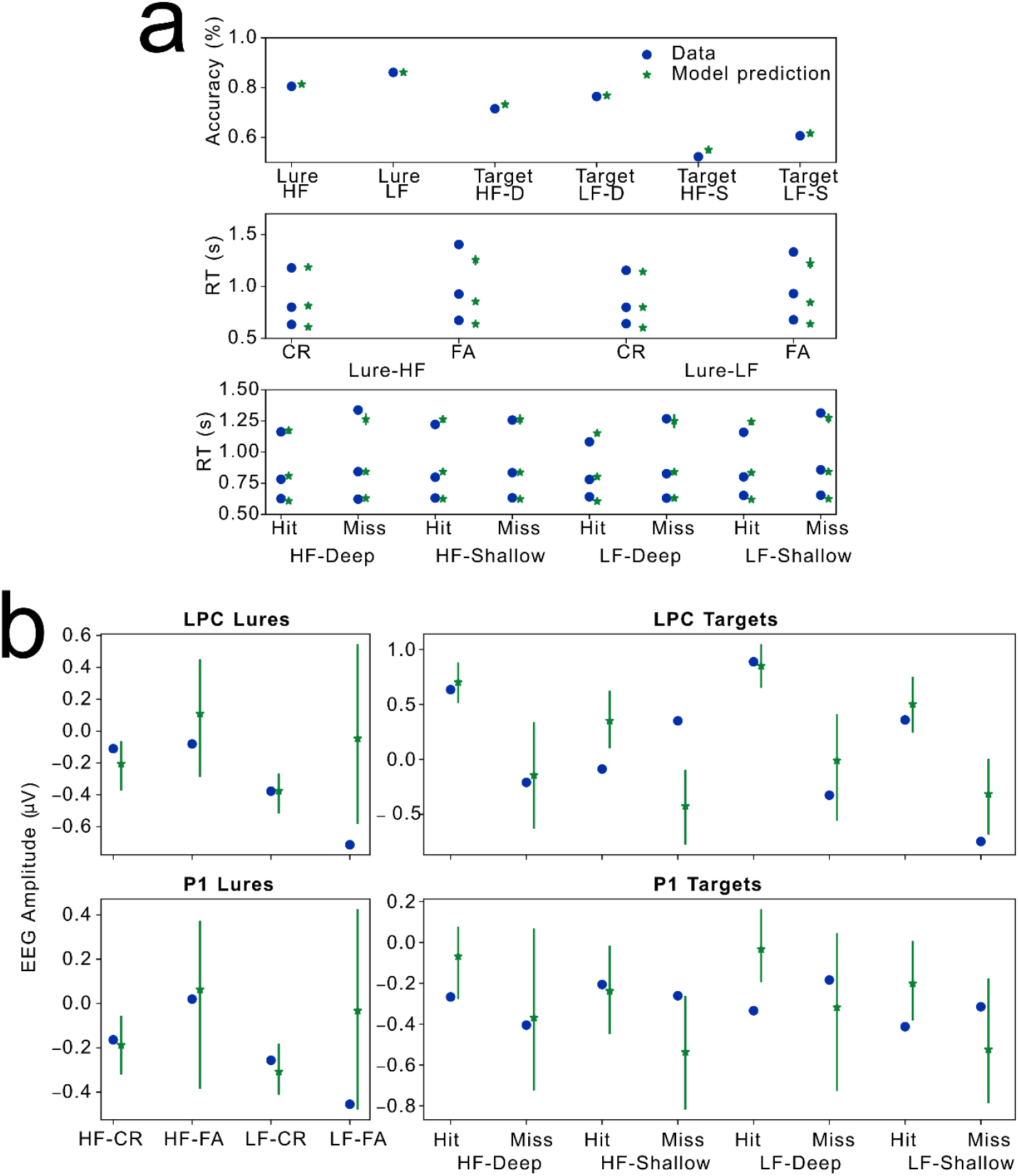
Model predictions for the model linking drift rate with LPC and visual P1 amplitudes. ***Note***. a. Model predictions of behavioural data based on the best fitting parameters across participants. b. Model predictions of mean LPC and P1visual P1 amplitudes based on the best fitting parameters across participants. The error band represents the 95% highest density interval across 200 samples generated from the best fitting parameters.

**Supplementary Figure 7.**
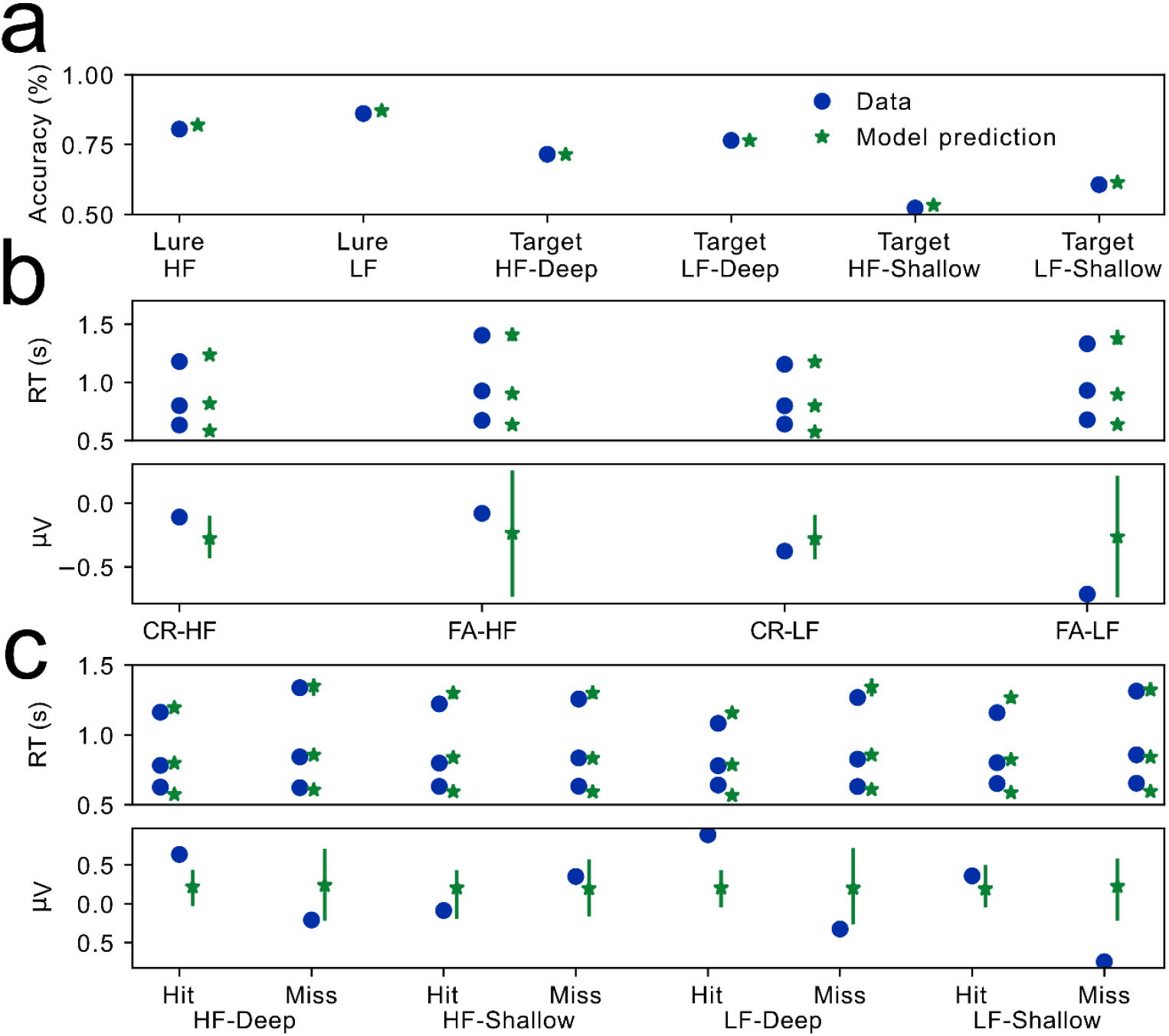
Model predictions for the model linking non-decision time with LPC amplitudes. ***Note*.** a. Model predictions of accuracy based on the best fitting parameters across participants. b. Model predictions of RTs and mean LPC amplitudes for lures. c. Model predictions of RTs and mean LPC amplitudes for targets. The error band represents the 95% highest density interval across 200 samples generated from the best fitting parameters.

**Supplementary Figure 8.**
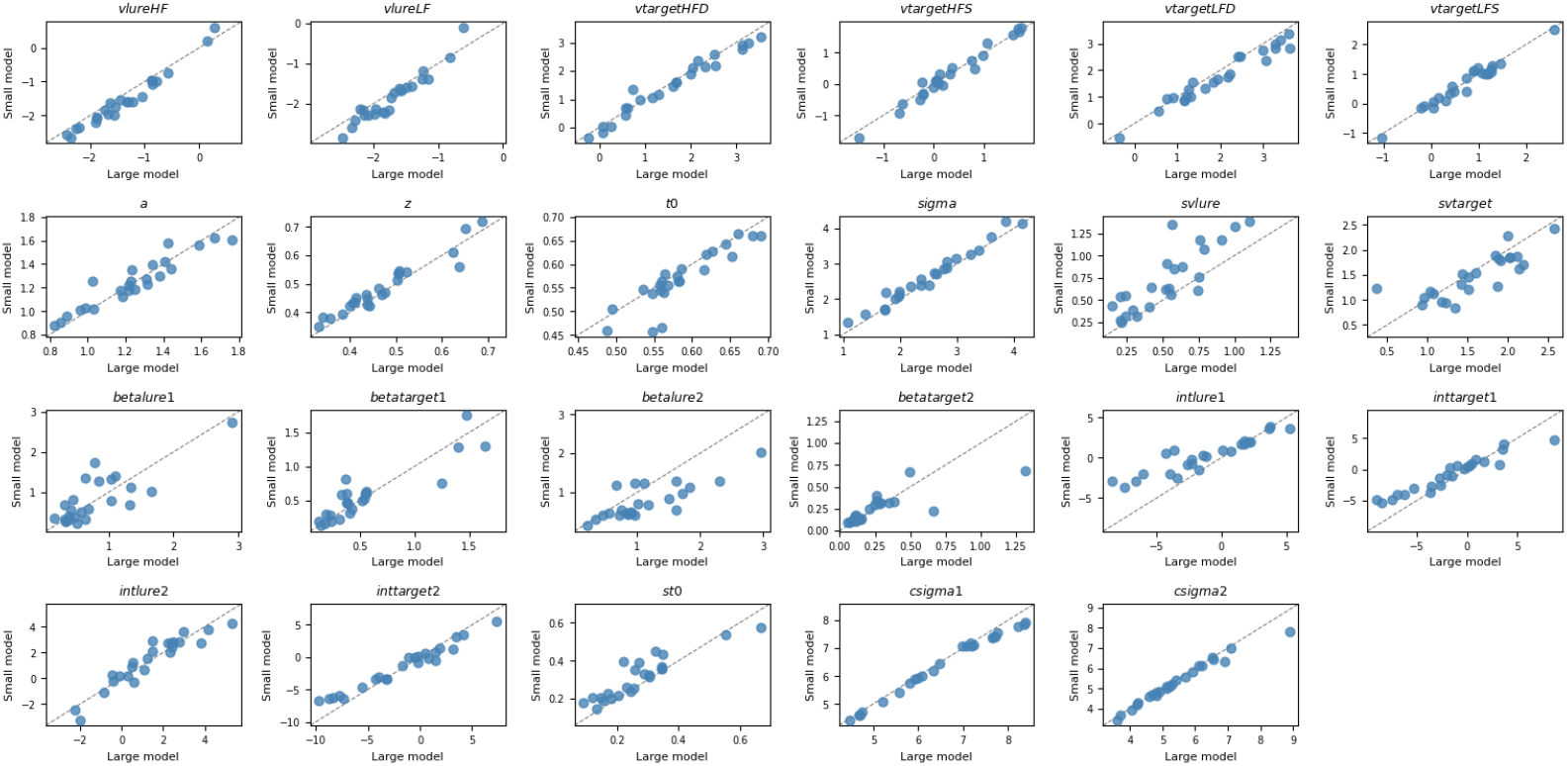
Comparisons of parameter estimates in Model 2 from models either trained on small or large number of trials per sample. ***Note***. The parameters were based on the median values from the posterior distributions after model fit to the experimental data. The small-sample model was trained on 300-500 trials per simulation; the large-sample model was trained on 1800-2400 trials per simulation.

**Supplementary Figure 9.**
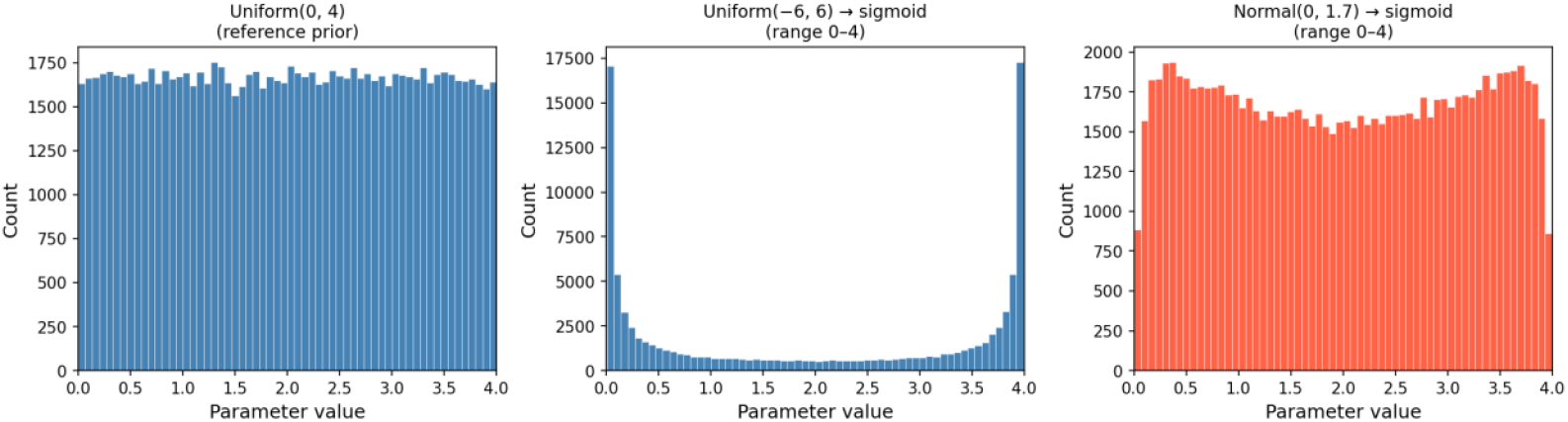
Examples of parameter distributions drawn from either a uniform distribution (left), a uniform distribution after transformation using a sigmoid function (middle), or a normal distribution after transformation using a sigmoid function (right).

**Supplementary Figure 10.**
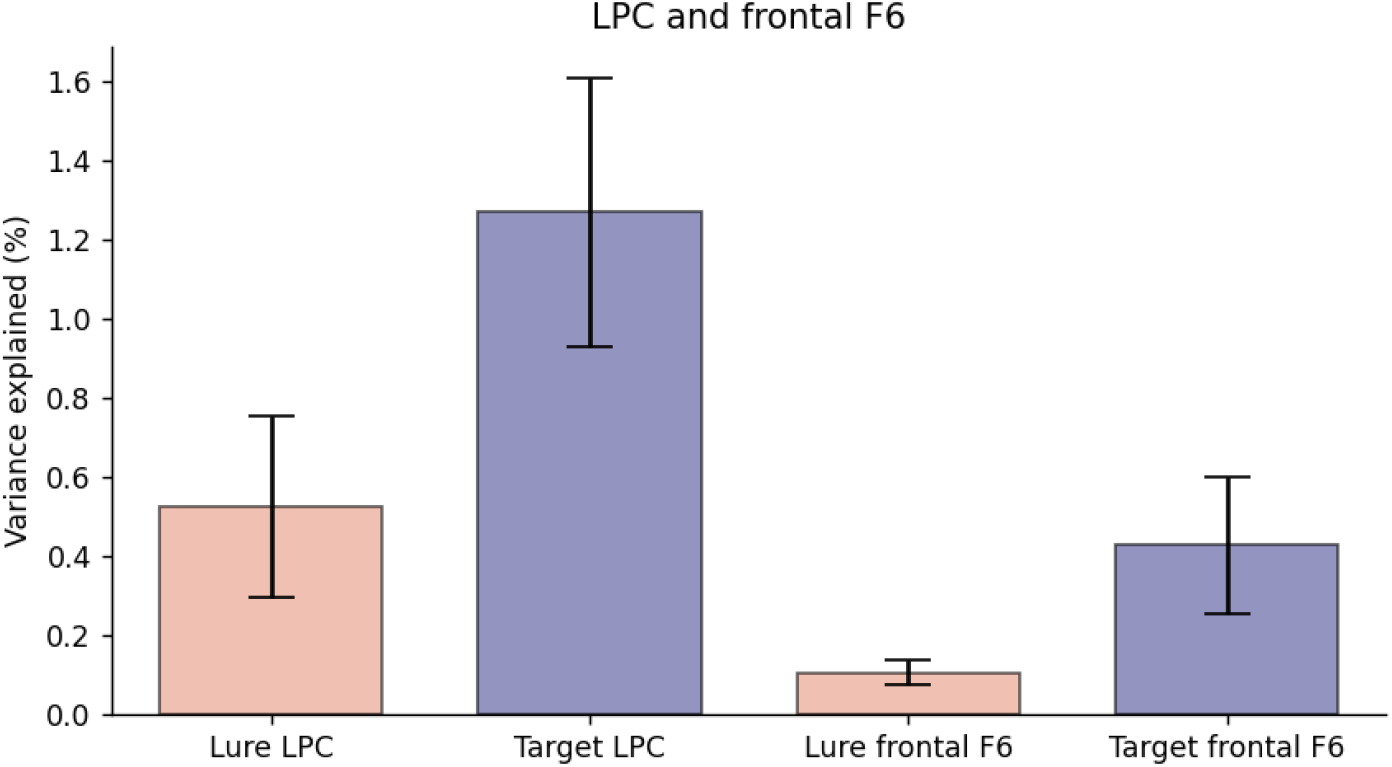
Variance-explained measures for the model that jointly predicts the LPC amplitude and also the pre-response amplitude at channel F6 after regressing out the LPC amplitude. Note. The error bar represents the 95% HDI.

**Supplementary Figure 11.**
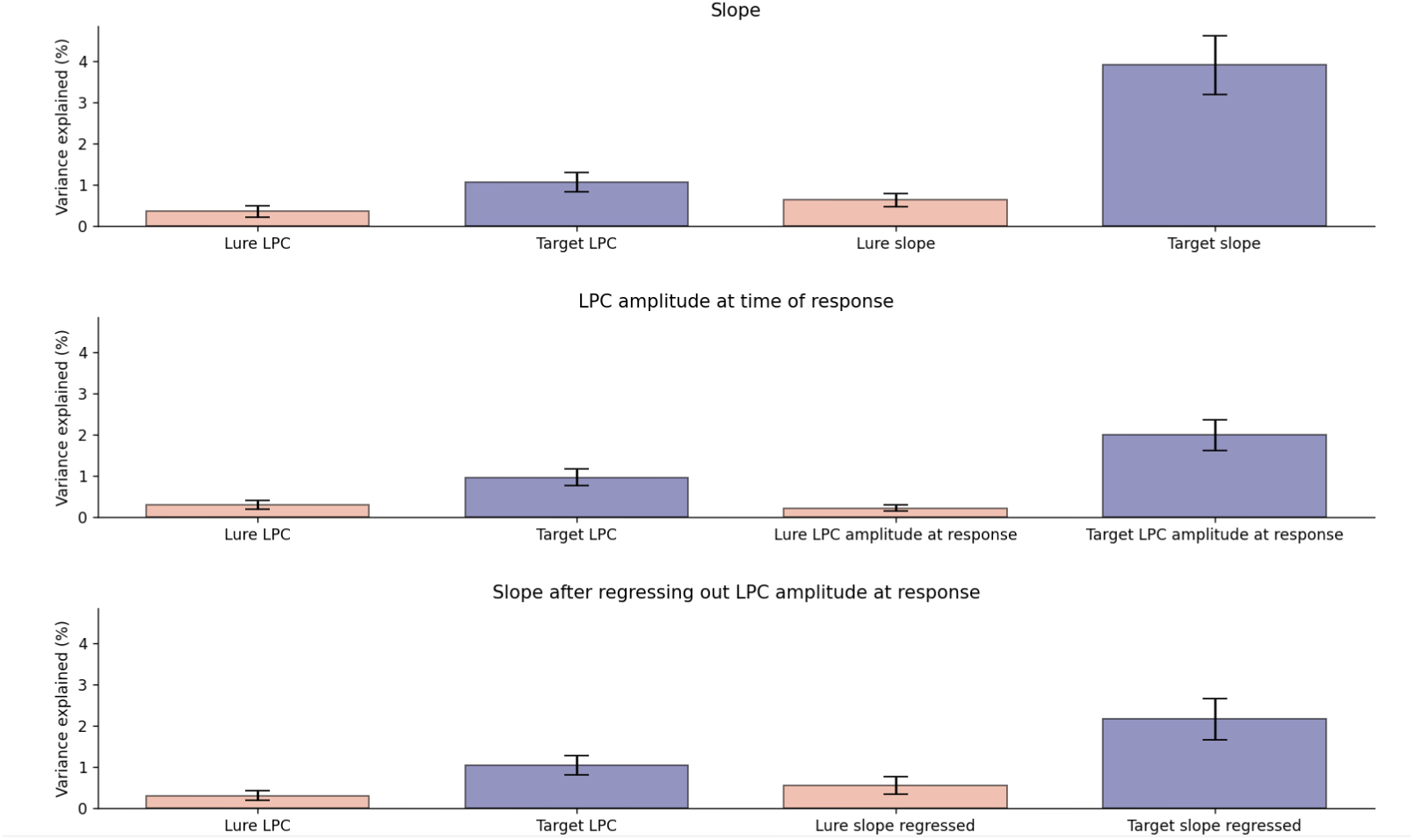
With the data after subtracting the stimulus-locked components, variance-explained measures for the model simultaneously predicts pre-response LPC amplitudes and LPC slopes estimated from -300 to 0 ms around the response. ***Note***. The model was set up the same way as the Model 2 but replacing prediction of P1 with the LPC slopes. The slope measures were estimated from -300 to 0 ms relative to the response and were rescaled to be consistent the range of predicted values of the neural component from initial training. The top panel shows the variance-explained measures across targets and lures for the prediction of the LPC amplitude and slope, separately. The middle panel shows the variance-explained measures across targets and lures for the prediction of the averaged pre-response LPC amplitude and the LPC amplitude at the time of the response, separately. The bottom panel shows the variance-explained measures across targets and lures for the prediction of the averaged pre-response LPC amplitude and residual of slope measures after regressing out the LPC amplitude at the time of response.

**Supplementary Table 1.**
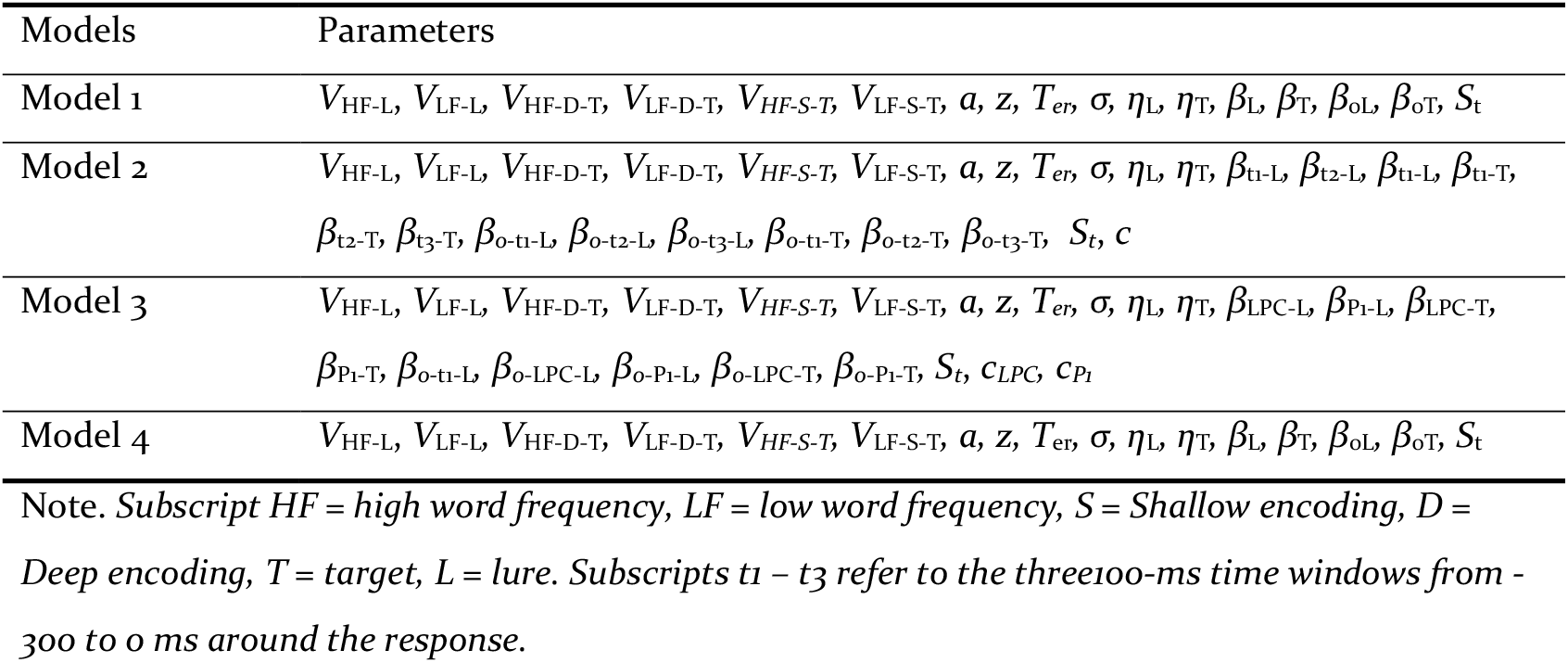
Parameters included in each model.

**Supplementary Table 2.**
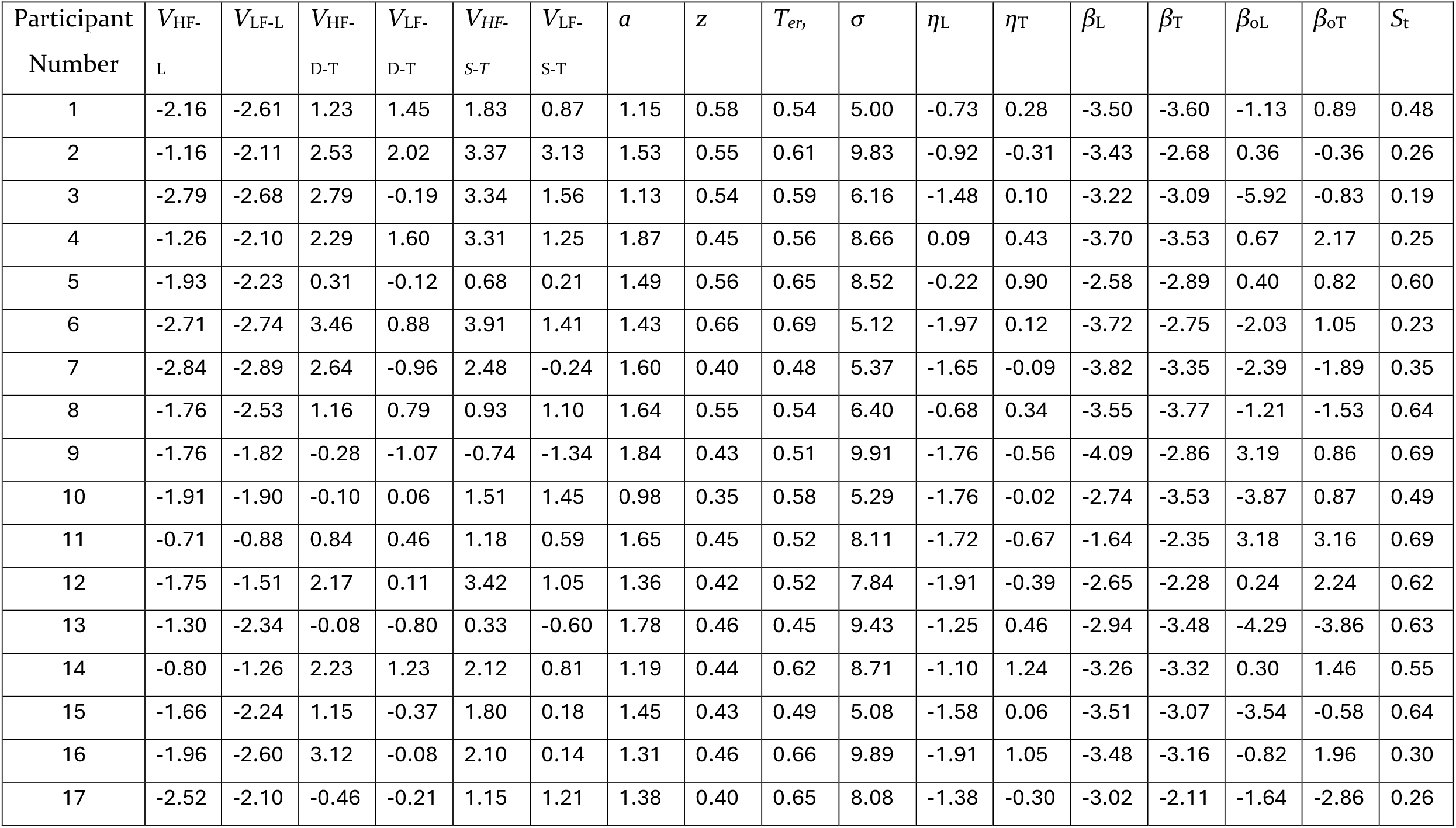
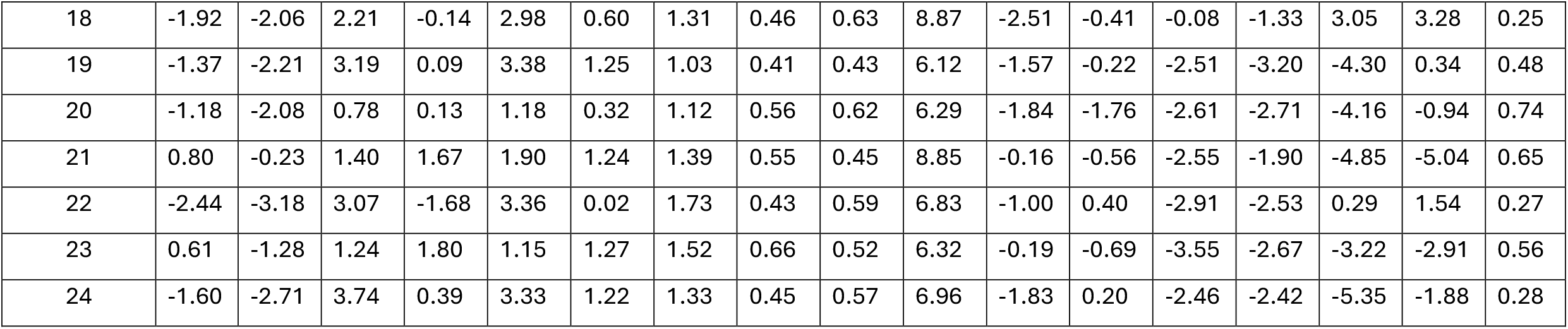
Model 1 median parameter estimates of the posterior distributions across participants.

| Participant<br>Number | $V_{HF-L}$ | $V_{LF-L}$ | $V_{HF-D-T}$ | $V_{LF-D-T}$ | $V_{HF-S-T}$ | $V_{LF-S-T}$ | $a$ | $z$ | $T_{er}$ | $\sigma$ | $\eta_L$ | $\eta_T$ | $\beta_L$ | $\beta_T$ | $\beta_{oL}$ | $\beta_{oT}$ | $S_t$ |
| --- | --- | --- | --- | --- | --- | --- | --- | --- | --- | --- | --- | --- | --- | --- | --- | --- | --- |
| 1 | -2.16 | -2.61 | 1.23 | 1.45 | 1.83 | 0.87 | 1.15 | 0.58 | 0.54 | 5.00 | -0.73 | 0.28 | -3.50 | -3.60 | -1.13 | 0.89 | 0.48 |
| 2 | -1.16 | -2.11 | 2.53 | 2.02 | 3.37 | 3.13 | 1.53 | 0.55 | 0.61 | 9.83 | -0.92 | -0.31 | -3.43 | -2.68 | 0.36 | -0.36 | 0.26 |
| 3 | -2.79 | -2.68 | 2.79 | -0.19 | 3.34 | 1.56 | 1.13 | 0.54 | 0.59 | 6.16 | -1.48 | 0.10 | -3.22 | -3.09 | -5.92 | -0.83 | 0.19 |
| 4 | -1.26 | -2.10 | 2.29 | 1.60 | 3.31 | 1.25 | 1.87 | 0.45 | 0.56 | 8.66 | 0.09 | 0.43 | -3.70 | -3.53 | 0.67 | 2.17 | 0.25 |
| 5 | -1.93 | -2.23 | 0.31 | -0.12 | 0.68 | 0.21 | 1.49 | 0.56 | 0.65 | 8.52 | -0.22 | 0.90 | -2.58 | -2.89 | 0.40 | 0.82 | 0.60 |
| 6 | -2.71 | -2.74 | 3.46 | 0.88 | 3.91 | 1.41 | 1.43 | 0.66 | 0.69 | 5.12 | -1.97 | 0.12 | -3.72 | -2.75 | -2.03 | 1.05 | 0.23 |
| 7 | -2.84 | -2.89 | 2.64 | -0.96 | 2.48 | -0.24 | 1.60 | 0.40 | 0.48 | 5.37 | -1.65 | -0.09 | -3.82 | -3.35 | -2.39 | -1.89 | 0.35 |
| 8 | -1.76 | -2.53 | 1.16 | 0.79 | 0.93 | 1.10 | 1.64 | 0.55 | 0.54 | 6.40 | -0.68 | 0.34 | -3.55 | -3.77 | -1.21 | -1.53 | 0.64 |
| 9 | -1.76 | -1.82 | -0.28 | -1.07 | -0.74 | -1.34 | 1.84 | 0.43 | 0.51 | 9.91 | -1.76 | -0.56 | -4.09 | -2.86 | 3.19 | 0.86 | 0.69 |
| 10 | -1.91 | -1.90 | -0.10 | 0.06 | 1.51 | 1.45 | 0.98 | 0.35 | 0.58 | 5.29 | -1.76 | -0.02 | -2.74 | -3.53 | -3.87 | 0.87 | 0.49 |
| 11 | -0.71 | -0.88 | 0.84 | 0.46 | 1.18 | 0.59 | 1.65 | 0.45 | 0.52 | 8.11 | -1.72 | -0.67 | -1.64 | -2.35 | 3.18 | 3.16 | 0.69 |
| 12 | -1.75 | -1.51 | 2.17 | 0.11 | 3.42 | 1.05 | 1.36 | 0.42 | 0.52 | 7.84 | -1.91 | -0.39 | -2.65 | -2.28 | 0.24 | 2.24 | 0.62 |
| 13 | -1.30 | -2.34 | -0.08 | -0.80 | 0.33 | -0.60 | 1.78 | 0.46 | 0.45 | 9.43 | -1.25 | 0.46 | -2.94 | -3.48 | -4.29 | -3.86 | 0.63 |
| 14 | -0.80 | -1.26 | 2.23 | 1.23 | 2.12 | 0.81 | 1.19 | 0.44 | 0.62 | 8.71 | -1.10 | 1.24 | -3.26 | -3.32 | 0.30 | 1.46 | 0.55 |
| 15 | -1.66 | -2.24 | 1.15 | -0.37 | 1.80 | 0.18 | 1.45 | 0.43 | 0.49 | 5.08 | -1.58 | 0.06 | -3.51 | -3.07 | -3.54 | -0.58 | 0.64 |
| 16 | -1.96 | -2.60 | 3.12 | -0.08 | 2.10 | 0.14 | 1.31 | 0.46 | 0.66 | 9.89 | -1.91 | 1.05 | -3.48 | -3.16 | -0.82 | 1.96 | 0.30 |
| 17 | -2.52 | -2.10 | -0.46 | -0.21 | 1.15 | 1.21 | 1.38 | 0.40 | 0.65 | 8.08 | -1.38 | -0.30 | -3.02 | -2.11 | -1.64 | -2.86 | 0.26 |
| 18 | -1.92 | -2.06 | 2.21 | -0.14 | 2.98 | 0.60 | 1.31 | 0.46 | 0.63 | 8.87 | -2.51 | -0.41 | -0.08 | -1.33 | 3.05 | 3.28 | 0.25 |
| 19 | -1.37 | -2.21 | 3.19 | 0.09 | 3.38 | 1.25 | 1.03 | 0.41 | 0.43 | 6.12 | -1.57 | -0.22 | -2.51 | -3.20 | -4.30 | 0.34 | 0.48 |
| 20 | -1.18 | -2.08 | 0.78 | 0.13 | 1.18 | 0.32 | 1.12 | 0.56 | 0.62 | 6.29 | -1.84 | -1.76 | -2.61 | -2.71 | -4.16 | -0.94 | 0.74 |
| 21 | 0.80 | -0.23 | 1.40 | 1.67 | 1.90 | 1.24 | 1.39 | 0.55 | 0.45 | 8.85 | -0.16 | -0.56 | -2.55 | -1.90 | -4.85 | -5.04 | 0.65 |
| 22 | -2.44 | -3.18 | 3.07 | -1.68 | 3.36 | 0.02 | 1.73 | 0.43 | 0.59 | 6.83 | -1.00 | 0.40 | -2.91 | -2.53 | 0.29 | 1.54 | 0.27 |
| 23 | 0.61 | -1.28 | 1.24 | 1.80 | 1.15 | 1.27 | 1.52 | 0.66 | 0.52 | 6.32 | -0.19 | -0.69 | -3.55 | -2.67 | -3.22 | -2.91 | 0.56 |
| 24 | -1.60 | -2.71 | 3.74 | 0.39 | 3.33 | 1.22 | 1.33 | 0.45 | 0.57 | 6.96 | -1.83 | 0.20 | -2.46 | -2.42 | -5.35 | -1.88 | 0.28 |

**Supplementary Table 3.**
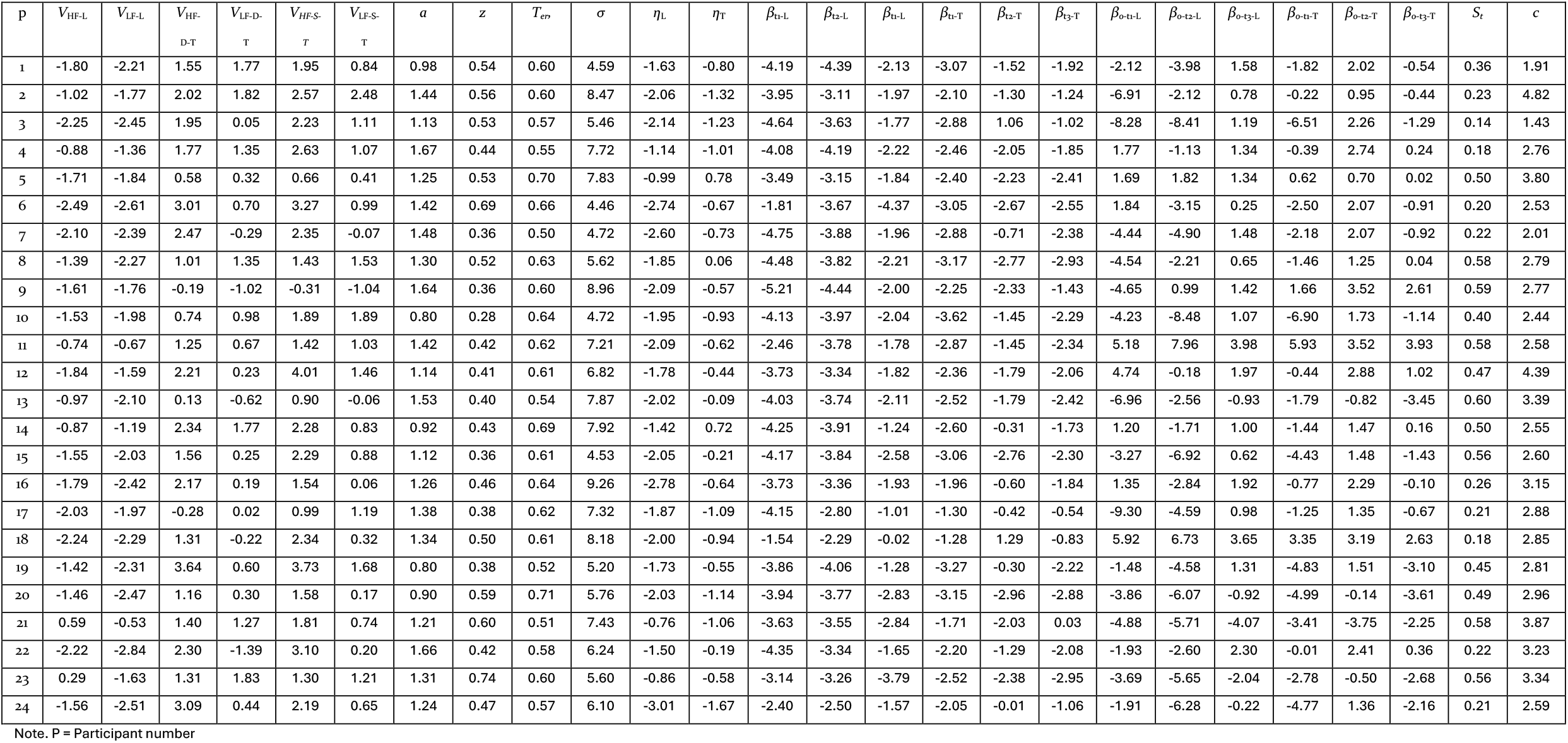
Model 2 median parameter estimates of the posterior distributions across participants.

**Supplementary Table 4.**
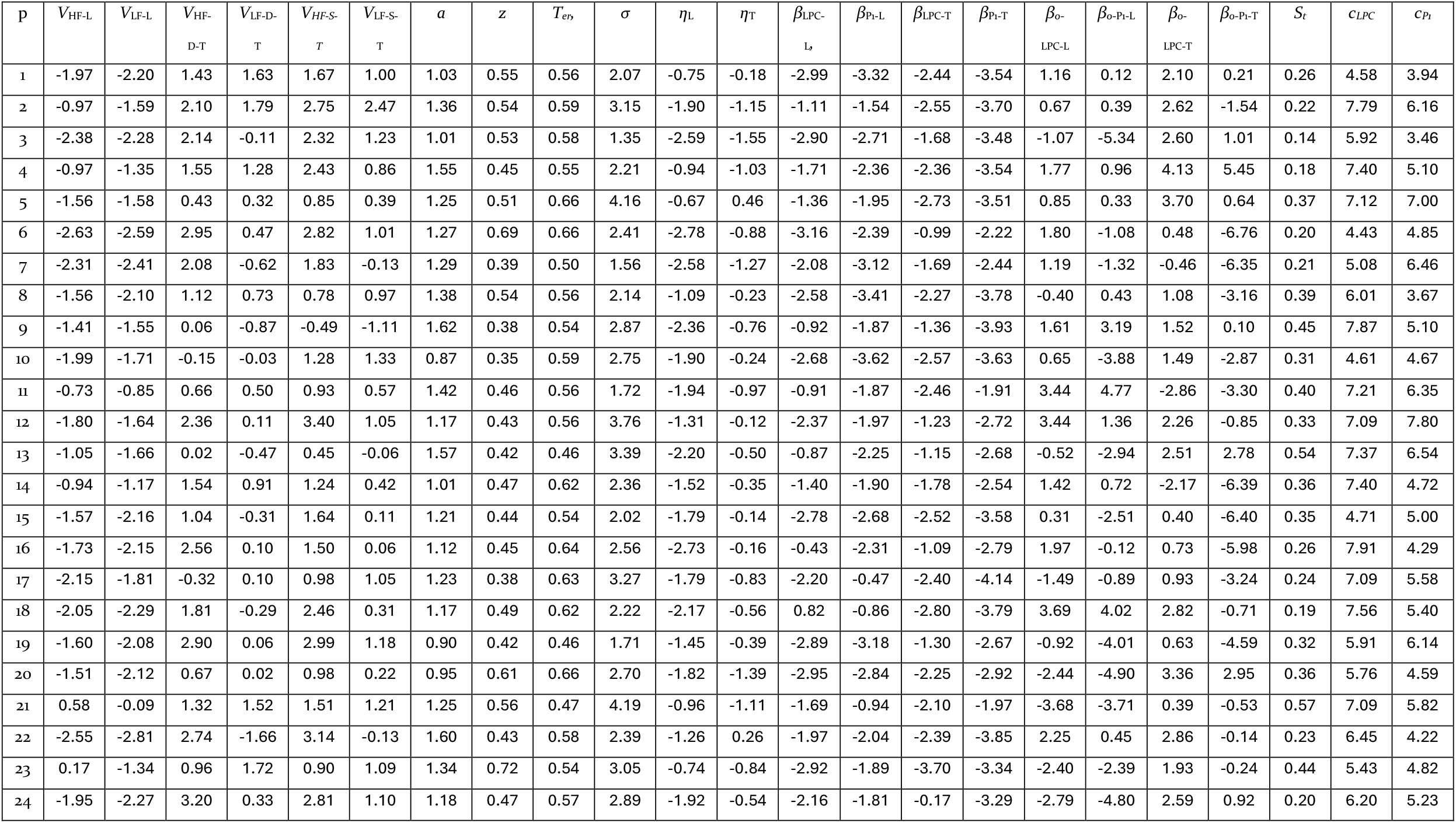
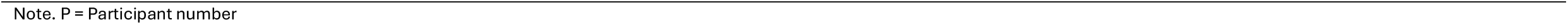
Model 3 median parameter estimates of the posterior distributions across participants.

**Supplementary Table 5.**
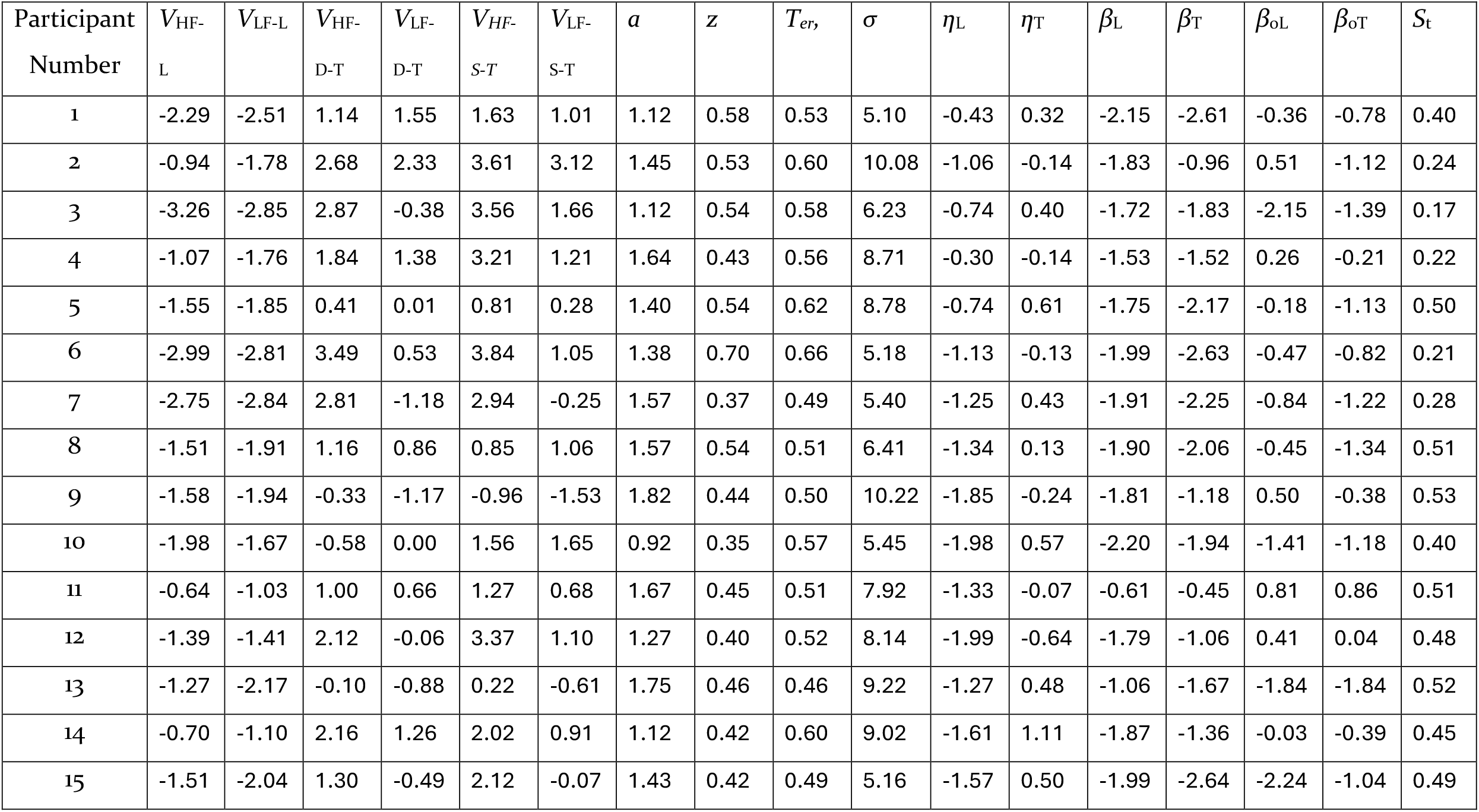
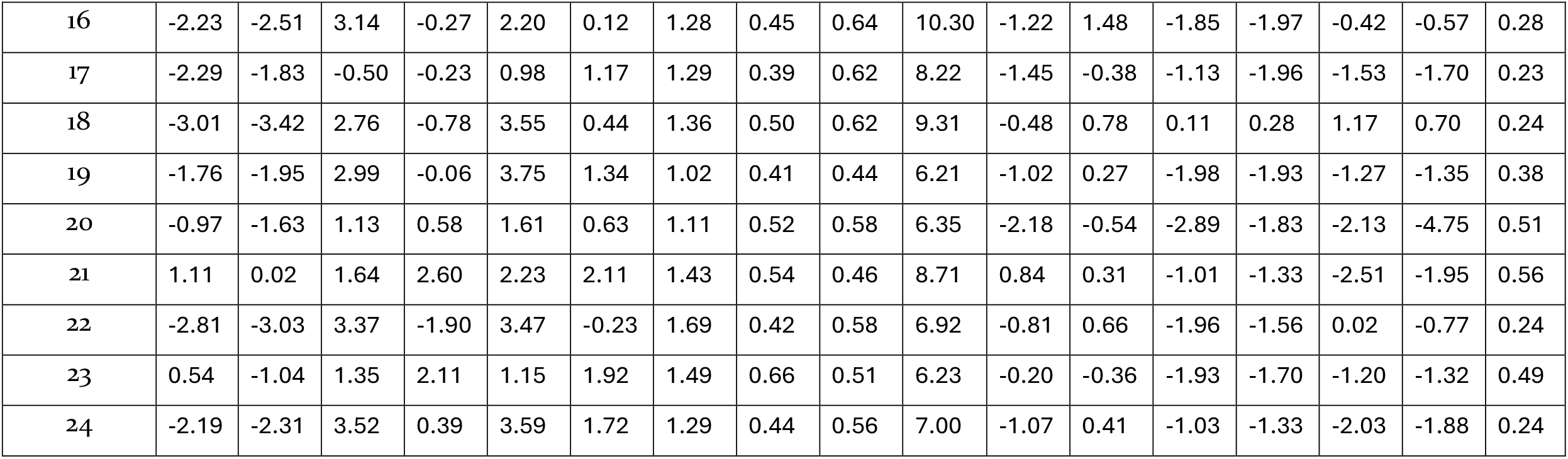
Model 4 median parameter estimates of the posterior distributions across participants.

| Participant<br>Number | $V_{HF-L}$ | $V_{LF-L}$ | $V_{HF-D-T}$ | $V_{LF-D-T}$ | $V_{HF-S-T}$ | $V_{LF-S-T}$ | $a$ | $z$ | $T_{er}$ | $\sigma$ | $\eta_L$ | $\eta_T$ | $\beta_L$ | $\beta_T$ | $\beta_{oL}$ | $\beta_{oT}$ | $S_t$ |
| --- | --- | --- | --- | --- | --- | --- | --- | --- | --- | --- | --- | --- | --- | --- | --- | --- | --- |
| 1 | -2.29 | -2.51 | 1.14 | 1.55 | 1.63 | 1.01 | 1.12 | 0.58 | 0.53 | 5.10 | -0.43 | 0.32 | -2.15 | -2.61 | -0.36 | -0.78 | 0.40 |
| 2 | -0.94 | -1.78 | 2.68 | 2.33 | 3.61 | 3.12 | 1.45 | 0.53 | 0.60 | 10.08 | -1.06 | -0.14 | -1.83 | -0.96 | 0.51 | -1.12 | 0.24 |
| 3 | -3.26 | -2.85 | 2.87 | -0.38 | 3.56 | 1.66 | 1.12 | 0.54 | 0.58 | 6.23 | -0.74 | 0.40 | -1.72 | -1.83 | -2.15 | -1.39 | 0.17 |
| 4 | -1.07 | -1.76 | 1.84 | 1.38 | 3.21 | 1.21 | 1.64 | 0.43 | 0.56 | 8.71 | -0.30 | -0.14 | -1.53 | -1.52 | 0.26 | -0.21 | 0.22 |
| 5 | -1.55 | -1.85 | 0.41 | 0.01 | 0.81 | 0.28 | 1.40 | 0.54 | 0.62 | 8.78 | -0.74 | 0.61 | -1.75 | -2.17 | -0.18 | -1.13 | 0.50 |
| 6 | -2.99 | -2.81 | 3.49 | 0.53 | 3.84 | 1.05 | 1.38 | 0.70 | 0.66 | 5.18 | -1.13 | -0.13 | -1.99 | -2.63 | -0.47 | -0.82 | 0.21 |
| 7 | -2.75 | -2.84 | 2.81 | -1.18 | 2.94 | -0.25 | 1.57 | 0.37 | 0.49 | 5.40 | -1.25 | 0.43 | -1.91 | -2.25 | -0.84 | -1.22 | 0.28 |
| 8 | -1.51 | -1.91 | 1.16 | 0.86 | 0.85 | 1.06 | 1.57 | 0.54 | 0.51 | 6.41 | -1.34 | 0.13 | -1.90 | -2.06 | -0.45 | -1.34 | 0.51 |
| 9 | -1.58 | -1.94 | -0.33 | -1.17 | -0.96 | -1.53 | 1.82 | 0.44 | 0.50 | 10.22 | -1.85 | -0.24 | -1.81 | -1.18 | 0.50 | -0.38 | 0.53 |
| 10 | -1.98 | -1.67 | -0.58 | 0.00 | 1.56 | 1.65 | 0.92 | 0.35 | 0.57 | 5.45 | -1.98 | 0.57 | -2.20 | -1.94 | -1.41 | -1.18 | 0.40 |
| 11 | -0.64 | -1.03 | 1.00 | 0.66 | 1.27 | 0.68 | 1.67 | 0.45 | 0.51 | 7.92 | -1.33 | -0.07 | -0.61 | -0.45 | 0.81 | 0.86 | 0.51 |
| 12 | -1.39 | -1.41 | 2.12 | -0.06 | 3.37 | 1.10 | 1.27 | 0.40 | 0.52 | 8.14 | -1.99 | -0.64 | -1.79 | -1.06 | 0.41 | 0.04 | 0.48 |
| 13 | -1.27 | -2.17 | -0.10 | -0.88 | 0.22 | -0.61 | 1.75 | 0.46 | 0.46 | 9.22 | -1.27 | 0.48 | -1.06 | -1.67 | -1.84 | -1.84 | 0.52 |
| 14 | -0.70 | -1.10 | 2.16 | 1.26 | 2.02 | 0.91 | 1.12 | 0.42 | 0.60 | 9.02 | -1.61 | 1.11 | -1.87 | -1.36 | -0.03 | -0.39 | 0.45 |
| 15 | -1.51 | -2.04 | 1.30 | -0.49 | 2.12 | -0.07 | 1.43 | 0.42 | 0.49 | 5.16 | -1.57 | 0.50 | -1.99 | -2.64 | -2.24 | -1.04 | 0.49 |
| 16 | -2.23 | -2.51 | 3.14 | -0.27 | 2.20 | 0.12 | 1.28 | 0.45 | 0.64 | 10.30 | -1.22 | 1.48 | -1.85 | -1.97 | -0.42 | -0.57 | 0.28 |
| 17 | -2.29 | -1.83 | -0.50 | -0.23 | 0.98 | 1.17 | 1.29 | 0.39 | 0.62 | 8.22 | -1.45 | -0.38 | -1.13 | -1.96 | -1.53 | -1.70 | 0.23 |
| 18 | -3.01 | -3.42 | 2.76 | -0.78 | 3.55 | 0.44 | 1.36 | 0.50 | 0.62 | 9.31 | -0.48 | 0.78 | 0.11 | 0.28 | 1.17 | 0.70 | 0.24 |
| 19 | -1.76 | -1.95 | 2.99 | -0.06 | 3.75 | 1.34 | 1.02 | 0.41 | 0.44 | 6.21 | -1.02 | 0.27 | -1.98 | -1.93 | -1.27 | -1.35 | 0.38 |
| 20 | -0.97 | -1.63 | 1.13 | 0.58 | 1.61 | 0.63 | 1.11 | 0.52 | 0.58 | 6.35 | -2.18 | -0.54 | -2.89 | -1.83 | -2.13 | -4.75 | 0.51 |
| 21 | 1.11 | 0.02 | 1.64 | 2.60 | 2.23 | 2.11 | 1.43 | 0.54 | 0.46 | 8.71 | 0.84 | 0.31 | -1.01 | -1.33 | -2.51 | -1.95 | 0.56 |
| 22 | -2.81 | -3.03 | 3.37 | -1.90 | 3.47 | -0.23 | 1.69 | 0.42 | 0.58 | 6.92 | -0.81 | 0.66 | -1.96 | -1.56 | 0.02 | -0.77 | 0.24 |
| 23 | 0.54 | -1.04 | 1.35 | 2.11 | 1.15 | 1.92 | 1.49 | 0.66 | 0.51 | 6.23 | -0.20 | -0.36 | -1.93 | -1.70 | -1.20 | -1.32 | 0.49 |
| 24 | -2.19 | -2.31 | 3.52 | 0.39 | 3.59 | 1.72 | 1.29 | 0.44 | 0.56 | 7.00 | -1.07 | 0.41 | -1.03 | -1.33 | -2.03 | -1.88 | 0.24 |

## References

Addante, R. J., Ranganath, C., & Yonelinas, A. P. (2012). Examining ERP correlates of recognition memory: Evidence of accurate source recognition without recollection. Neuroimage, 62(1), 439–450.

Ally, B. A., Simons, J. S., McKeever, J. D., Peers, P. V., & Budson, A. E. (2008). Parietal contributions to recollection: electrophysiological evidence from aging and patients with parietal lesions. Neuropsychologia, 46(7), 1800–1812.

Bainbridge, W. A. (2017). The memorability of people: Intrinsic memorability across transformations of a person’s face. Journal of Experimental Psychology: Learning, Memory, and Cognition, 43(5), 706.

Bode, S., Feuerriegel, D., Bennett, D., & Alday, P. M. (2019). The Decision Decoding ToolBOX (DDTBOX)–A multivariate pattern analysis toolbox for event-related potentials. Neuroinformatics, 17, 27–42.

Brezis, N., Bronfman, Z. Z., Yovel, G., & Goshen-Gottstein, Y. (2017). The electrophysiological signature of remember–know is confounded with memory strength and cannot be interpreted as evidence for dual-process theory of recognition. Journal of cognitive neuroscience, 29(2), 322–336.

Bridger, E. K., Bader, R., & Mecklinger, A. (2014). More ways than one: ERPs reveal multiple familiarity signals in the word frequency mirror effect. Neuropsychologia, 57, 179–190.

Brockdorff, N., & Lamberts, K. (2000). A feature-sampling account of the time course of old–new recognition judgments. Journal of Experimental Psychology: Learning, Memory, and Cognition, 26(1), 77.

Brosnan, M. B., Sabaroedin, K., Silk, T., Genc, S., Newman, D. P., Loughnane, G. M., Fornito, A., O’Connell, R. G., & Bellgrove, M. A. (2020). Evidence accumulation during perceptual decisions in humans varies as a function of dorsal frontoparietal organization. Nature Human Behaviour, 4(8), 844–855.

Cassey, P. J., Gaut, G., Steyvers, M., & Brown, S. D. (2016). A generative joint model for spike trains and saccades during perceptual decision-making. Psychonomic bulletin & review, 23, 1757–1778.

Chang, M., Johns, B. T., & Brainerd, C. J. (2025). True and false recognition in MINERVA2: Integrating fuzzy-trace theory and computational memory modeling. Psychological review.

Chen, H., Heathcote, A., Sauer, J. D., Palmer, M. A., & Osth, A. F. (2024). Greater target or lure variability? An exploration on the effects of stimulus types and memory paradigms. Memory & Cognition, 52(3), 554–573.

Cox, G. E., & Shiffrin, R. M. (2017). A dynamic approach to recognition memory. Psychological review, 124(6), 795.

Craik, F. I., & Lockhart, R. S. (1972). Levels of processing: A framework for memory research. Journal of Verbal Learning and Verbal Behavior, 11(6), 671–684.

Delorme, A., & Makeig, S. (2004). EEGLAB: an open source toolbox for analysis of single-trial EEG dynamics including independent component analysis. J Neurosci Methods, 134(1), 9–21. 10.1016/j.jneumeth.2003.10.009

Desender, K., Donner, T. H., & Verguts, T. (2021). Dynamic expressions of confidence within an evidence accumulation framework. Cognition, 207, 104522.

Di Russo, F., Martínez, A., Sereno, M. I., Pitzalis, S., & Hillyard, S. A. (2002). Cortical sources of the early components of the visual evoked potential. Human brain mapping, 15(2), 95–111.

Dobbins, I. G., Rice, H. J., Wagner, A. D., & Schacter, D. L. (2003). Memory orientation and success: separable neurocognitive components underlying episodic recognition. Neuropsychologia, 41(3), 318–333.

Donaldson, D. I., Wheeler, M. E., & Petersen, S. E. (2010). Remember the source: dissociating frontal and parietal contributions to episodic memory. Journal of cognitive neuroscience, 22(2), 377–391.

Egan, J. P. (1958). Recognition memory and the operating characteristic. USAF Operational Applications Laboratory Technical Note.

Fengler, A., Govindarajan, L. N., Chen, T., & Frank, M. J. (2021). Likelihood approximation networks (LANs) for fast inference of simulation models in cognitive neuroscience. Elife, 10, e65074.

Feuerriegel, D., Jiwa, M., Turner, W. F., Andrejević, M., Hester, R., & Bode, S. (2021). Tracking dynamic adjustments to decision making and performance monitoring processes in conflict tasks. Neuroimage, 238, 118265.

Feuerriegel, D., Murphy, M., Konski, A., Mepani, V., Sun, J., Hester, R., & Bode, S. (2022). Electrophysiological correlates of confidence differ across correct and erroneous perceptual decisions. Neuroimage, 259, 119447.

Fong, L. C., Garrett, P. M., Smith, P. L., Hester, R., Bode, S., & Feuerriegel, D. (2025). Tracing the neural trajectories of evidence accumulation and motor preparation processes during voluntary decisions. BioRxiv, 2025.2010. 2030.685712.

Forstmann, B. U., Dutilh, G., Brown, S., Neumann, J., Von Cramon, D. Y., Ridderinkhof, K. R., & Wagenmakers, E.-J. (2008). Striatum and pre-SMA facilitate decision-making under time pressure. Proceedings of the National Academy of Sciences, 105(45), 17538–17542.

Frank, M. J., Gagne, C., Nyhus, E., Masters, S., Wiecki, T. V., Cavanagh, J. F., & Badre, D. (2015). fMRI and EEG predictors of dynamic decision parameters during human reinforcement learning. Journal of neuroscience, 35(2), 485–494.

Friedman, D., & Johnson, R. (2000). Event-related potential (ERP) studies of memory encoding and retrieval: A selective review. Microscopy research and technique, 51(1), 6–28.

Frömer, R., Nassar, M. R., Ehinger, B. V., & Shenhav, A. (2024). Common neural choice signals can emerge artefactually amid multiple distinct value signals. Nature Human Behaviour, 8(11), 2194–2208.

Fukuda, K., Printzlau, F. A., Kang, M.-S., & Woodman, G. F. (2024). Posterior ERP tracks evidence accumulation for memory-based decisions. (pre-print).

Ghaderi-Kangavari, A., Rad, J. A., & Nunez, M. D. (2023). A general integrative neurocognitive modeling framework to jointly describe EEG and decision-making on single trials. Computational Brain & Behavior, 6(3), 317–376.

Grogan, J. P., Rys, W., Kelly, S. P., & O’Connell, R. G. (2023). Confidence is predicted by pre-and post-choice decision signal dynamics. Imaging Neuroscience, 1, 1–23.

Groppe, D. M., Urbach, T. P., & Kutas, M. (2011). Mass univariate analysis of event-related brain potentials/fields I: A critical tutorial review. Psychophysiology, 48(12), 1711–1725.

Guo, C., Zhu, Y., Ding, J., Fan, S., & Paller, K. A. (2004). An electrophysiological investigation of memory encoding, depth of processing, and word frequency in humans. Neuroscience letters, 356(2), 79–82.

Harris, J. D., Cutmore, T. R., O’Gorman, J., Finnigan, S., & Shum, D. H. (2013). Electrophysiological correlates of perceptual auditory priming without explicit recognition memory. Journal of psychophysiology.

Hebart, M. N., Zheng, C. Y., Pereira, F., & Baker, C. I. (2020). Revealing the multidimensional mental representations of natural objects underlying human similarity judgements. Nature Human Behaviour, 4(11), 1173–1185.

Henson, R. N., Rugg, M., Shallice, T., Josephs, O., & Dolan, R. J. (1999). Recollection and familiarity in recognition memory: an event-related functional magnetic resonance imaging study. Journal of neuroscience, 19(10), 3962–3972.

Hower, K. H., Wixted, J., Berryhill, M. E., & Olson, I. R. (2014). Impaired perception of mnemonic oldness, but not mnemonic newness, after parietal lobe damage. Neuropsychologia, 56, 409–417.

Hutchinson, J. B., Uncapher, M. R., & Wagner, A. D. (2015). Increased functional connectivity between dorsal posterior parietal and ventral occipitotemporal cortex during uncertain memory decisions. Neurobiology of learning and memory, 117, 71–83.

Johnson, R., Pfefferbaum, A., & Kopell, B. S. (1985). P300 and long-term memory: Latency predicts recognition performance. Psychophysiology, 22(5), 497–507.

Jung, T.-P., Makeig, S., Humphries, C., Lee, T.-W., Mckeown, M. J., Iragui, V., & Sejnowski, T. J. (2000). Removing electroencephalographic artifacts by blind source separation. Psychophysiology, 37(2), 163–178.

Kahn, I., Davachi, L., & Wagner, A. D. (2004). Functional-neuroanatomic correlates of recollection: implications for models of recognition memory. Journal of neuroscience, 24(17), 4172–4180.

Kelly, S. P., Corbett, E. A., & O’Connell, R. G. (2021). Neurocomputational mechanisms of prior-informed perceptual decision-making in humans. Nature Human Behaviour, 5(4), 467–481.

Kelly, S. P., & O’Connell, R. G. (2013). Internal and external influences on the rate of sensory evidence accumulation in the human brain. Journal of neuroscience, 33(50), 19434–19441.

Kim, M. S., Kwon, J. S., Kang, S. S., Youn, T., & Kang, K. W. (2004). Impairment of recognition memory in schizophrenia: Event-related potential study using a continuous recognition task. Psychiatry and clinical neurosciences, 58(5), 465–472.

Ko, Y. H., Zhou, A., Niessen, E., Stahl, J., Weiss, P. H., Hester, R., Bode, S., & Feuerriegel, D. (2024). Neural correlates of confidence during decision formation in a perceptual judgment task. Cortex, 173, 248–262.

Kriegeskorte, N., Simmons, W. K., Bellgowan, P. S., & Baker, C. I. (2009). Circular analysis in systems neuroscience: the dangers of double dipping. Nature neuroscience, 12(5), 535–540.

Kwon, S., Rugg, M. D., Wiegand, R., Curran, T., & Morcom, A. M. (2023). A meta-analysis of event-related potential correlates of recognition memory. Psychonomic bulletin & review, 30(6), 2083–2105.

Lenzi, A., Bessac, J., Rudi, J., & Stein, M. L. (2023). Neural networks for parameter estimation in intractable models. Computational Statistics & Data Analysis, 185, 107762.

Leynes, P. A., & Mok, B. A. (2017). Encoding focus alters diagnostic recollection and event-related potentials (ERPs). Brain and cognition, 117, 1–11.

MacLeod, C. M., & Kampe, K. E. (1996). Word frequency effects on recall, recognition, and word fragment completion tests. Journal of Experimental Psychology: Learning, Memory, and Cognition, 22(1), 132.

Malmberg, K. J., & Nelson, T. O. (2003). The word frequency effect for recognition memory and the elevated-attention hypothesis. Memory & Cognition, 31(1), 35–43.

Maris, E., & Oostenveld, R. (2007). Nonparametric statistical testing of EEG-and MEG-data. Journal of neuroscience methods, 164(1), 177–190. 10.1016/j.jneumeth.2007.03.024

Morey, R. D. (2008). Confidence intervals from normalized data: A correction to Cousineau (2005). Tutorials in quantitative methods for psychology, 4(2), 61–64.

Nie, A., Pan, R., & Shen, H. (2021). How processing fluency contributes to the old/new effects of familiarity and recollection: Evidence from the remember/know paradigm. The American journal of psychology, 134(3), 297–319.

Nosofsky, R. M., Ehinger, K. A., & Osth, A. F. (2025). Tests of a hybrid-similarity exemplar model of context-dependent memorability in a high-dimensional real-world category domain. Journal of Experimental Psychology: General.

Nunez, M. D., Vandekerckhove, J., & Srinivasan, R. (2017). How attention influences perceptual decision making: Single-trial EEG correlates of drift-diffusion model parameters. Journal of Mathematical Psychology, 76, 117–130.

O’connell, R. G., Dockree, P. M., & Kelly, S. P. (2012a). A supramodal accumulation-to-bound signal that determines perceptual decisions in humans. Nature neuroscience, 15(12), 1729–1735.

O’connell, R. G., Dockree, P. M., & Kelly, S. P. (2012b). A supramodal accumulation-to-bound signal that determines perceptual decisions in humans. Nature neuroscience, 15(12), 1729. 10.1038/nn.3248

O’Connell, R. G., & Kelly, S. P. (2021). Neurophysiology of human perceptual decision-making. Annual review of neuroscience, 44(1), 495–516.

O’Connell, R. G., Parés-Pujolràs, E., Corbett, E. A., Feuerriegel, D., & Kelly, S. P. (2025). Regressing away common neural choice signals does not make them artifacts: Comment on Frömer et al. 2024. Imaging Neuroscience, 3, IMAG. a. 60.

Olichney, J., Taylor, J., Gatherwright, J., Salmon, D., Bressler, A., Kutas, M., & Iragui-Madoz, V. (2008). Patients with MCI and N400 or P600 abnormalities are at very high risk for conversion to dementia. Neurology, 70(19_part_2), 1763–1770.

Onyper, S. V., Zhang, Y. X., & Howard, M. W. (2010). Some-or-none recollection: Evidence from item and source memory. Journal of Experimental Psychology: General, 139(2), 341.

Osth, A. F., Bora, B., Dennis, S., & Heathcote, A. (2017). Diffusion vs. linear ballistic accumulation: Different models, different conclusions about the slope of the zROC in recognition memory. Journal of Memory and Language, 96, 36–61.

Osth, A. F., & Dennis, S. (2020). Global matching models of recognition memory.

Osth, A. F., & Zhang, L. (2024). Integrating word-form representations with global similarity computation in recognition memory. Psychonomic bulletin & review, 31(3), 1000–1031.

Osth, A. F., Zhang, L., & Williams, S. (2025). A global matching model of choice and response times in the Deese–Roediger–Mcdermott semantic and structural false recognition paradigms. Psychological review.

Osth, A. F., Zhou, J., Chen, H., & Sun, J. (2025). Sequential sampling models in memory. Learning & memory: A comprehensive reference, 107–126.

Ouyang, G., Herzmann, G., Zhou, C., & Sommer, W. (2011). Residue iteration decomposition (RIDE): A new method to separate ERP components on the basis of latency variability in single trials. Psychophysiology, 48(12), 1631–1647.

Ouyang, G., Hildebrandt, A., Sommer, W., & Zhou, C. (2017). Exploiting the intra-subject latency variability from single-trial event-related potentials in the P3 time range: a review and comparative evaluation of methods. Neuroscience & Biobehavioral Reviews, 75, 1–21.

Ouyang, G., Sommer, W., & Zhou, C. (2015). A toolbox for residue iteration decomposition (RIDE)—A method for the decomposition, reconstruction, and single trial analysis of event related potentials. Journal of neuroscience methods, 250, 7–21.

Peirce, J. W. (2007). PsychoPy—psychophysics software in Python. Journal of neuroscience methods, 162(1-2), 8–13.

Philiastides, M. G., Heekeren, H. R., & Sajda, P. (2014). Human scalp potentials reflect a mixture of decision-related signals during perceptual choices. Journal of neuroscience, 34(50), 16877–16889.

Pisauro, M. A., Fouragnan, E., Retzler, C., & Philiastides, M. G. (2017). Neural correlates of evidence accumulation during value-based decisions revealed via simultaneous EEG-fMRI. Nature communications, 8(1), 15808.

Pleskac, T. J., & Busemeyer, J. R. (2010). Two-stage dynamic signal detection: a theory of choice, decision time, and confidence. Psychological review, 117(3), 864.

Radev, S. T., D’Alessandro, M., Mertens, U. K., Voss, A., Köthe, U., & Bürkner, P.-C. (2021). Amortized bayesian model comparison with evidential deep learning. IEEE transactions on neural networks and learning systems, 34(8), 4903–4917.

Radev, S. T., Graw, F., Chen, S., Mutters, N. T., Eichel, V. M., Bärnighausen, T., & Köthe, U. (2021). OutbreakFlow: Model-based Bayesian inference of disease outbreak dynamics with invertible neural networks and its application to the COVID-19 pandemics in Germany. PLoS computational biology, 17(10), e1009472.

Radev, S. T., Mertens, U. K., Voss, A., Ardizzone, L., & Köthe, U. (2020). BayesFlow: Learning complex stochastic models with invertible neural networks. IEEE transactions on neural networks and learning systems, 33(4), 1452–1466.

Ratcliff, R. (1978). A theory of memory retrieval. Psychological review, 85(2), 59.

Ratcliff, R., Gomez, P., & McKoon, G. (2004). A diffusion model account of the lexical decision task. Psychological review, 111(1), 159.

Ratcliff, R., McKoon, G., & Tindall, M. (1994). Empirical generality of data from recognition memory receiver-operating characteristic functions and implications for the global memory models. Journal of Experimental Psychology: Learning, Memory, and Cognition, 20(4), 763.

Ratcliff, R., Sederberg, P. B., Smith, T. A., & Childers, R. (2016). A single trial analysis of EEG in recognition memory: Tracking the neural correlates of memory strength. Neuropsychologia, 93, 128–141.

Ratcliff, R., & Tuerlinckx, F. (2002). Estimating parameters of the diffusion model: Approaches to dealing with contaminant reaction times and parameter variability. Psychonomic bulletin & review, 9(3), 438–481.

Reid, J. N., Guitard, D., & Jamieson, R. K. (2026). MINERVA OPS: A computational framework for the representation and recognition of orthographic, phonological, and semantic associates. Psychonomic bulletin & review, 33(2), 77.

Reid, J. N., & Jamieson, R. K. (2023). True and false recognition in MINERVA 2: Extension to sentences and metaphors. Journal of Memory and Language, 129, 104397.

Rubin, S. R., Petten, C. V., Glisky, E. L., & Newberg, W. M. (1999). Memory conjunction errors in younger and older adults: Event-related potential and neuropsychological data. Cognitive neuropsychology, 16(3-5), 459–488.

Rugg, M. D. (1990). Event-related brain potentials dissociate repetition effects of high- and low-frequency words. Memory & Cognition, 18(4), 367–379.

Rugg, M. D., Cox, C. J., Doyle, M. C., & Wells, T. (1995). Event-related potentials and the recollection of low and high frequency words. Neuropsychologia, 33(4), 471–484.

Rugg, M. D., & Curran, T. (2007). Event-related potentials and recognition memory. Trends in cognitive sciences, 11(6), 251–257.

Rugg, M. D., Mark, R. E., Walla, P., Schloerscheidt, A. M., Birch, C. S., & Allan, K. (1998). Dissociation of the neural correlates of implicit and explicit memory. Nature, 392(6676), 595–598.

Rugg, M. D., & Yonelinas, A. P. (2003). Human recognition memory: a cognitive neuroscience perspective. Trends in cognitive sciences, 7(7), 313–319.

Rutishauser, U., Aflalo, T., Rosario, E. R., Pouratian, N., & Andersen, R. A. (2018). Single-neuron representation of memory strength and recognition confidence in left human posterior parietal cortex. Neuron, 97(1), 209–220. e203.

Sainsbury-Dale, M., Zammit-Mangion, A., & Huser, R. (2024). Likelihood-free parameter estimation with neural Bayes estimators. The American Statistician, 78(1), 1–14.

Sarah, S. Y., & Rugg, M. D. (2010). Dissociation of the electrophysiological correlates of familiarity strength and item repetition. Brain research, 1320, 74–84.

Schmitt, M., Bürkner, P.-C., Köthe, U., & Radev, S. T. (2023). Detecting model misspecification in amortized Bayesian inference with neural networks. DAGM German Conference on Pattern Recognition,

Sestieri, C., Shulman, G. L., & Corbetta, M. (2017). The contribution of the human posterior parietal cortex to episodic memory. Nature Reviews Neuroscience, 18(3), 183–192.

Sestieri, C., Tosoni, A., Mignogna, V., McAvoy, M. P., Shulman, G. L., Corbetta, M., & Romani, G. L. (2014). Memory accumulation mechanisms in human cortex are independent of motor intentions. Journal of neuroscience, 34(20), 6993–7006.

Smith, N. K., Cacioppo, J. T., Larsen, J. T., & Chartrand, T. L. (2003). May I have your attention, please: Electrocortical responses to positive and negative stimuli. Neuropsychologia, 41(2), 171–183.

Smith, P. L., & Lilburn, S. D. (2020). Vision for the blind: Visual psychophysics and blinded inference for decision models. Psychonomic bulletin & review, 27(5), 882–910.

Smith, P. L., & Ratcliff, R. (2009). An integrated theory of attention and decision making in visual signal detection. Psychological review, 116(2), 283.

Sokratous, K., Fitch, A. K., & Kvam, P. D. (2023). How to ask twenty questions and win: Machine learning tools for assessing preferences from small samples of willingness-to-pay prices. Journal of choice modelling, 48, 100418.

Starns, J. J., & Ratcliff, R. (2014). Validating the unequal-variance assumption in recognition memory using response time distributions instead of ROC functions: A diffusion model analysis. Journal of Memory and Language, 70, 36–52.

Steinemann, N. A., O’Connell, R. G., & Kelly, S. P. (2018). Decisions are expedited through multiple neural adjustments spanning the sensorimotor hierarchy. Nature communications, 9(1), 3627.

Sun, J., Feuerriegel, D., & Osth, A. F. (2025). Can Systematic Drift Rate Variability Replace Random Variability in the Diffusion Decision Model?

Sun, J., Osth, A. F., & Feuerriegel, D. (2024). The late positive event-related potential component is time locked to the decision in recognition memory tasks. Cortex, 176, 194–208.

Tarhan, L., De Freitas, J., & Konkle, T. (2021). Behavioral and neural representations en route to intuitive action understanding. Neuropsychologia, 163, 108048.

Turner, B. M., Forstmann, B. U., Wagenmakers, E.-J., Brown, S. D., Sederberg, P. B., & Steyvers, M. (2013). A Bayesian framework for simultaneously modeling neural and behavioral data. Neuroimage, 72, 193–206.

van Vugt, M. K., Beulen, M. A., & Taatgen, N. A. (2019). Relation between centro-parietal positivity and diffusion model parameters in both perceptual and memory-based decision making. Brain research, 1715, 1–12.

Vilberg, K. L., & Rugg, M. D. (2008). Memory retrieval and the parietal cortex: a review of evidence from a dual-process perspective. Neuropsychologia, 46(7), 1787–1799.

Vilberg, K. L., & Rugg, M. D. (2009). Functional significance of retrieval-related activity in lateral parietal cortex: Evidence from fMRI and ERPs. Human brain mapping, 30(5), 1490–1501.

von Krause, M., Radev, S. T., & Voss, A. (2022). Mental speed is high until age 60 as revealed by analysis of over a million participants. Nature Human Behaviour, 6(5), 700–708.

Voss, J. L., & Paller, K. A. (2009). Remembering and knowing: Electrophysiological distinctions at encoding but not retrieval. Neuroimage, 46(1), 280–289.

Wagner, A. D., Shannon, B. J., Kahn, I., & Buckner, R. L. (2005). Parietal lobe contributions to episodic memory retrieval. Trends in cognitive sciences, 9(9), 445–453.

Wheeler, M. E., & Buckner, R. L. (2003). Functional dissociation among components of remembering: control, perceived oldness, and content. Journal of neuroscience, 23(9), 3869–3880.

Wheeler, M. E., & Buckner, R. L. (2004). Functional-anatomic correlates of remembering and knowing. Neuroimage, 21(4), 1337–1349.

Widmann, A., Schröger, E., & Maess, B. (2015). Digital filter design for electrophysiological data–a practical approach. Journal of neuroscience methods, 250, 34–46.

Wieschen, E. M., Makani, A., Radev, S. T., Voss, A., & Spaniol, J. (2024). Age-related differences in decision-making: Evidence accumulation is more gradual in older age. Experimental Aging Research, 50(5), 537–549.

Wixted, J. T. (2007). Dual-process theory and signal-detection theory of recognition memory. Psychological review, 114(1), 152.

Wixted, J. T., & Mickes, L. (2010). A continuous dual-process model of remember/know judgments. Psychological review, 117(4), 1025.

Wolk, D. A., Manning, K., Kliot, D., & Arnold, S. E. (2013). Recognition memory in amnestic-mild cognitive impairment: insights from event-related potentials. Frontiers in Aging Neuroscience, 5, 89.

Woodruff, C. C., Hayama, H. R., & Rugg, M. D. (2006). Electrophysiological dissociation of the neural correlates of recollection and familiarity. Brain research, 1100(1), 125–135.

Woroch, B., & Gonsalves, B. D. (2010). Event-related potential correlates of item and source memory strength. Brain research, 1317, 180–191.

Yang, H., Laforge, G., Stojanoski, B., Nichols, E. S., McRae, K., & Köhler, S. (2019). Late positive complex in event-related potentials tracks memory signals when they are decision relevant. Scientific reports, 9(1), 9469.

Ye, J., Nie, A., & Liu, S. (2019). How do word frequency and memory task influence directed forgetting: An ERP study. International Journal of Psychophysiology, 146, 157–172.

Yonelinas, A. P. (2002). The nature of recollection and familiarity: A review of 30 years of research. Journal of Memory and Language, 46(3), 441–517.

Yu, S. S., Johnson, J. D., & Rugg, M. D. (2012). Dissociation of recollection-related neural activity in ventral lateral parietal cortex. Cognitive neuroscience, 3(3-4), 142–149.

Zammit-Mangion, A., Sainsbury-Dale, M., & Huser, R. (2024). Neural methods for amortized inference. Annual Review of Statistics and Its Application, 12.

Zhang, L., & Osth, A. F. (2024). Modelling orthographic similarity effects in recognition memory reveals support for open bigram representations of letter coding. Cognitive psychology, 148, 101619.

Zhang, S., & Sutton, R. S. (2017). A deeper look at experience replay. arXiv preprint arXiv:1712.01275.

